# Lysosomal Enhancement Prevents Infection with PrP^Sc^, α-Synuclein & Tau Prions

**DOI:** 10.1101/2025.06.24.661349

**Authors:** Robert C.C. Mercer, Nhat T.T. Le, Nadia A. Mirza-Romero, Erin Flaherty, Joseph P. DeFranco, Giada Lavigna, Isabel C. Orbe, Jean R. P. Gatdula, Douglas G. Fraser, Aravind Sundaravadivelu, Janelle S. Vultaggio, Aaron B. Beeler, Roberto Chiesa, Glenn C. Telling, David A. Harris

## Abstract

Prion diseases are fatal neurodegenerative diseases of humans and other mammals with no current treatment options. Here, we describe the characterization of a novel anti-prion compound, elacridar (GW120918), which has sub-micromolar activity in assays of prion infection, propagation and toxicity. Elacridar acts at an early step in the prion infection process, enhancing degradation of newly formed PrP^Sc^. The lysosome is the likely site of elacridar’s anti-prion effects, based on transcriptomic analysis and the use of functional lysosomal probes. Elacridar alters gene expression networks controlling lysosomal sterol and lipid metabolism but, unlike other lysosomotropic drugs, it prominently upregulates genes that control lysosomal pH. Surprisingly, these effects occur independently of TFEB nuclear translocation, suggesting novel regulatory mechanisms. The anti-prion effects of elacridar extend to α-synuclein and tau prions, highlighting lysosomal enhancement as a general strategy for treatment of protein misfolding neurodegenerative diseases.

**Graphical abstract:** 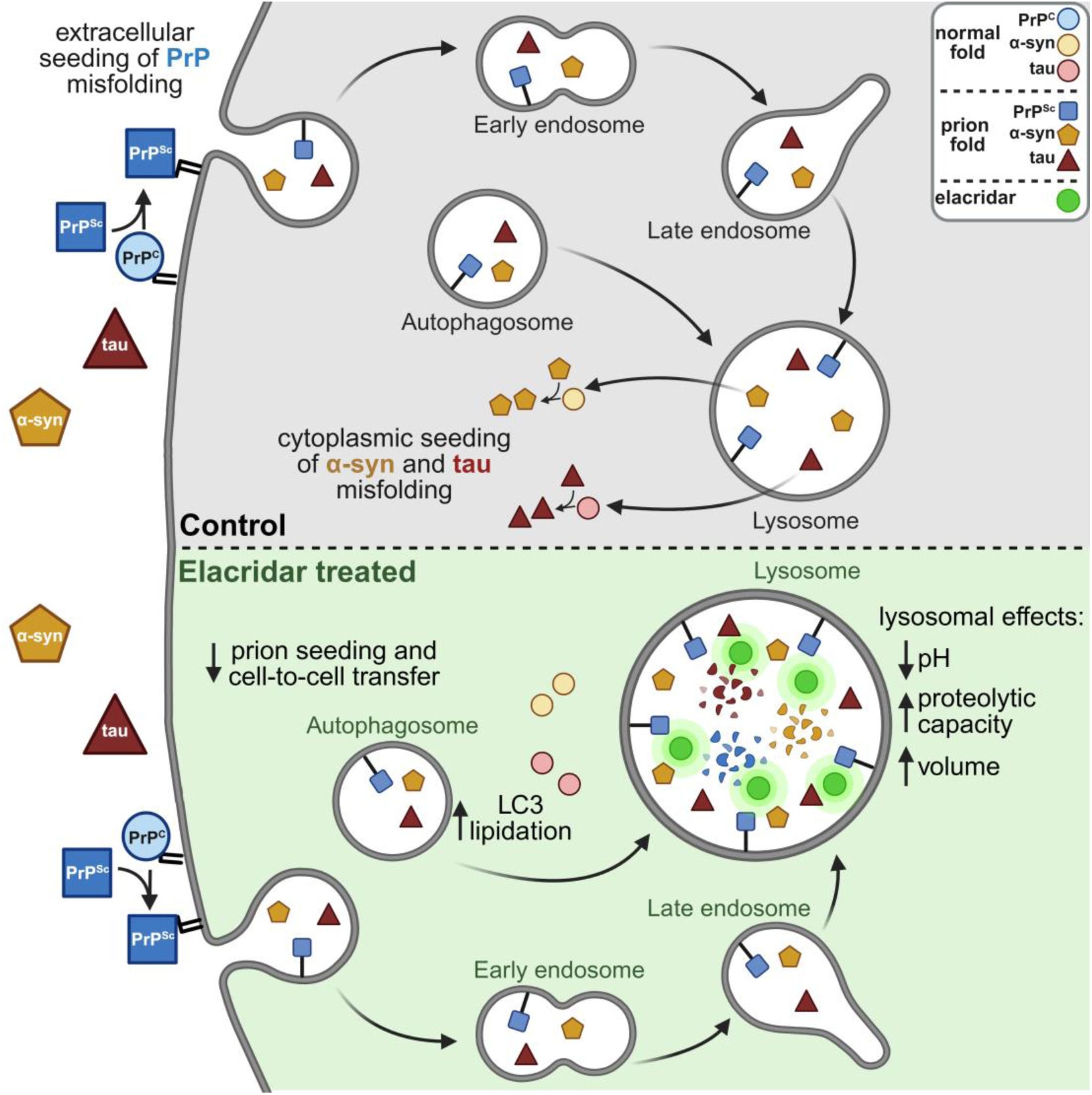

## Introduction

Prion diseases are invariably fatal neurodegenerative diseases of humans and other animals for which there are no available treatment options (1). These diseases are driven by the structural conversion of the primarily α-helical prion protein (PrP^C^) to a pathogenic conformation (PrP^Sc^), which forms fibrils of parallel in-register intermolecular β-sheets (2) and is capable of propagation via a self-templating mechanism (3).

The quest to identify small molecules for the treatment of prion disease has been ongoing for more than 30 years without success (4). Numerous compounds have been identified that can prevent prion propagation in cell culture, but many fail in animal models, and none have shown efficacy in patients (5,6). Several factors have contributed to this failure, including the confounding effect of cell division on determinations of compound efficacy *in vitro* (7–9), strain specific responses to small molecule therapeutics (10–12), and the lack of a clear molecular target for many anti-prion compounds (13,14).

While studying the anti-prion effects of ligands of the sigma receptors, σ1R and σ2R (14), we were led to examine elacridar (GW120918), a third-generation inhibitor of the xenobiotic transporter MDR1/p-glycoprotein/ABCB1, due to its reported interaction with σ2R (15). Surprisingly, we found that elacridar is a potent anti-prion compound with sub-micromolar efficacy using *in vitro* assays of prion infection, propagation and toxicity. Subsequent experimentation revealed that elacridar acts through a unique mechanism of lysosomal activation and that its anti-prion effects extend to pathological protein aggregates of α-synuclein and tau, ultimately suggesting new therapeutic strategies for all protein misfolding neurodegenerative diseases.

## Results

### Elacridar is a potent anti-prion compound

In a previous study, we reported that several synthetic ligands of the sigma receptors (σ1R/*Sigmar1* and σ2R/*Tmem97*) inhibit prion propagation and neurotoxicity *in vitro* (14). Surprisingly, we found that these anti-prion effects were independent of σR receptor binding. Moreover, because there was no discernable effect of these compounds on total or cell-surface levels of PrP^C^ or on prion seeding activity, we hypothesized that their action involves other cellular targets or pathways (14). During the course of these experiments, we examined elacridar, a previously identified σ2R ligand and clinically used inhibitor of the MDR-1 drug efflux transporter (15–17).

We confirmed the high-affinity interaction of elacridar with σ2R (15) using the National Institute of Mental Health Psychoactive Drug Screening Program (PDSP), which assays the binding affinity between small molecules and a variety of CNS receptors and transporters (18). We found that elacridar also binds to σ1R with 100x lower affinity than σ2R (ki σ2R = 24 nM; ki σ1R = 2615 nM; Supplementary Table 1). We then examined the effect of elacridar on prion propagation using multiple cell types (N2a, L929, CAD5) chronically infected with multiple murine prion strains (RML, 22L, ME7), and determined EC_50_ values for reducing the levels of proteinase K (PK)-resistant PrP (hereafter referred to operationally as PrP^Sc^) ranging from 298 to 926 nM (Figure 1A, Table 1, Supplementary Figure 1). Long-term elacridar treatment of N2a cells chronically infected with RML prions permanently cured them of infection, even after the compound was removed (Figure 1B). Elacridar was also efficacious against RML and sheep SSBP/1 prions using the scrapie cell assay (19,20) with RK13 cells expressing murine and ovine PrPs (21,22) (Supplementary Figure 2).

**Figure 1:**
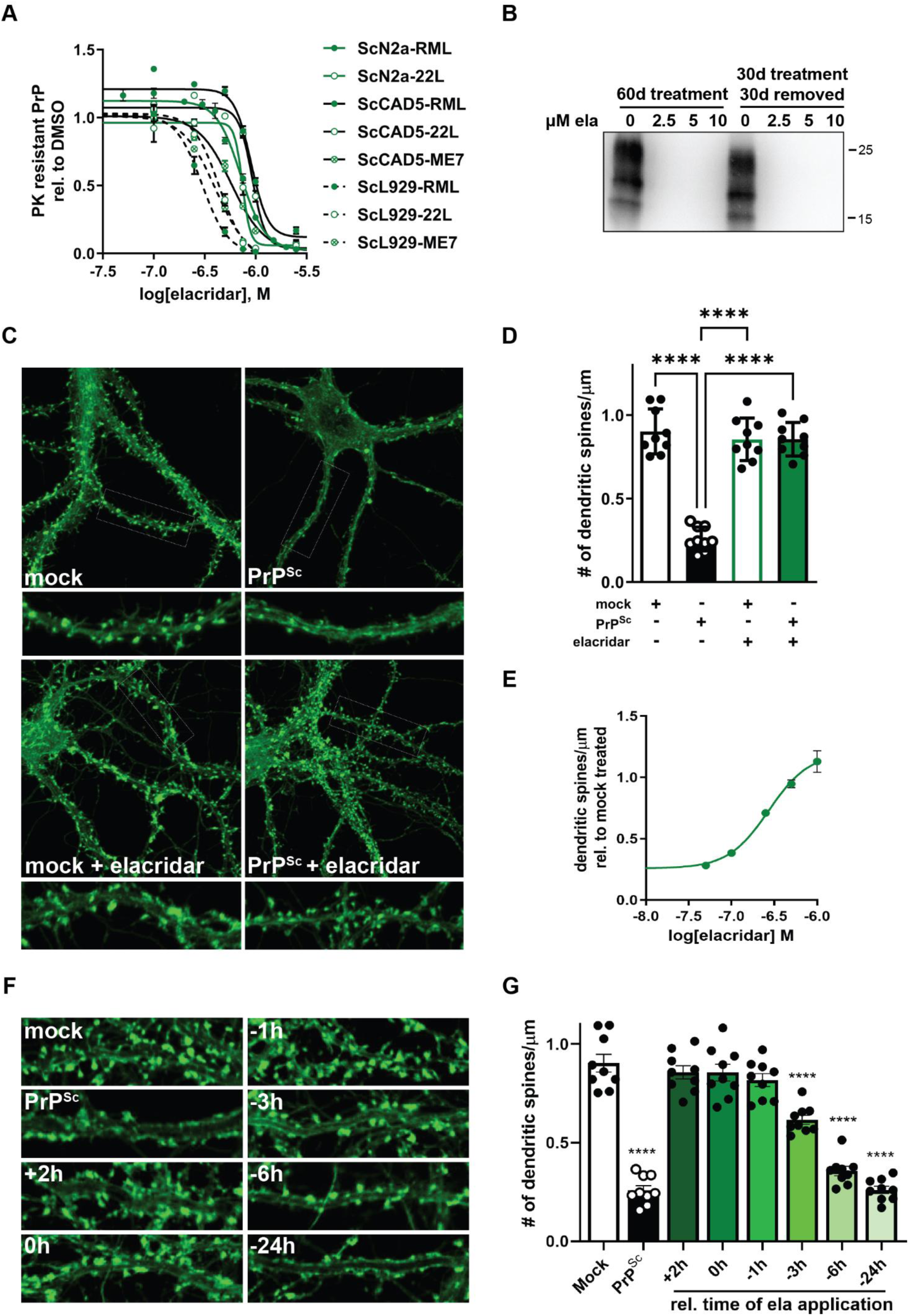
Anti-prion activity of elacridar in cell lines and neurons. **A)** Dose response curves for the effect of elacridar on the level of PK-resistant PrP in N2a, CAD5, and L929 cells chronically infected (Sc) with RML, 22L, or ME7 prions. Cells were treated for 7 days prior to western blotting. All data points represent three independent replicates. **B)** Elacridar permanently cures RML-infected N2a cells. Cells were cultured in the presence of the indicated concentration of elacridar for 60 days, or for 30 days with elacridar followed by 30 days in DMSO vehicle. Western blot analysis reveals an absence of PK-resistant PrP following either treatment. **C)** Elacridar prevents prion-induced collapse of hippocampal dendritic spines. Primary hippocampal neuronal cultures were exposed to 500 nM elacridar for 2 h before application of PrP^Sc^ purified from RML-infected brains (PrP^Sc^), or mock purified material from uninfected brains (mock). F-actin was imaged using Alexa Fluor 488 conjugated phalloidin to visualize dendritic spines. **D)** Mean number of dendritic spines/μm ± SEM presented in **C** is plotted. **E)** Elacridar prevents prion-induced dendritic spine collapse in a dose dependent manner. Primary hippocampal neuronal cultures were exposed to increasing concentrations of elacridar before application of purified RML prions. The number of dendritic spines remaining after 24h is plotted relative to mock treated cultures. **F)** 500 nM elacridar does not protect hippocampal dendritic spines if application if delayed more than 1 hour following prion exposure. Time of elacridar addition after exposure of neurons to purified RML prions is indicated in hours. F-actin was imaged using Alexa Fluor 488 conjugated phalloidin to visualize dendritic spines. **G)** Mean number of dendritic spines/μm ± SEM shown in **F** is plotted. For all quantifications, p < 0.0001 = ****; p < 0.001 = ***; p < 0.01 = **; p < 0.05 = *; ns = not significant using one-way ANOVA and Dunnett’s multiple comparisons test.

**Table 1:**
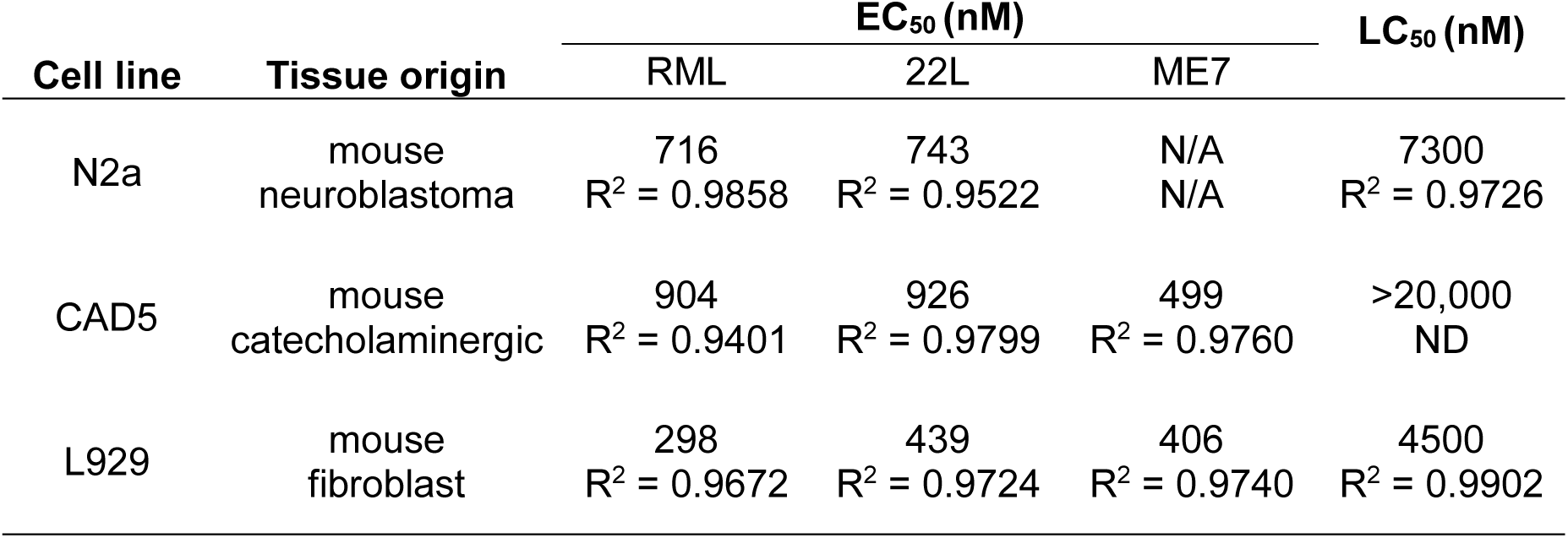
Anti-prion properties of elacridar in immortalized cells.

| Cell line | Tissue origin | EC<sub>50</sub> (nM) |  |  | LC<sub>50</sub> (nM) |
| --- | --- | --- | --- | --- | --- |
|  |  | RML | 22L | ME7 |  |
| N2a | mouse neuroblastoma | 716<br>R<sup>2</sup> = 0.9858 | 743<br>R<sup>2</sup> = 0.9522 | N/A<br>N/A | 7300<br>R<sup>2</sup> = 0.9726 |
| CAD5 | mouse catecholaminergic | 904<br>R<sup>2</sup> = 0.9401 | 926<br>R<sup>2</sup> = 0.9799 | 499<br>R<sup>2</sup> = 0.9760 | >20,000<br>ND |
| L929 | mouse fibroblast | 298<br>R<sup>2</sup> = 0.9672 | 439<br>R<sup>2</sup> = 0.9724 | 406<br>R<sup>2</sup> = 0.9740 | 4500<br>R<sup>2</sup> = 0.9902 |

We next examined the effect of elacridar on prion-mediated neurotoxicity using a hippocampal neuron dendritic spine retraction assay that we have employed extensively in the investigation of other anti-prion compounds (14,23–27). Exposure of neurons to highly enriched preparations of PrP^Sc^ (28) induced a rapid PrP^C^ dependent retraction of dendritic spines (26), an effect that was completely prevented by 500 nM elacridar with an EC_50_ of 272 nM (Figure 1C-E). Interestingly, a delay in the application of elacridar relative to RML prions by of little as 3 hours resulted in a time-dependent decrease in elacridar’s protective effect (Figure 1F, G). This result suggested that elacridar acts on cellular processes downstream of the initial conversion of PrP^C^ to PrP^Sc^, which has been reported to occur on the cell surface within minutes of prion exposure (29,30).

### Elacridar does not directly inhibit the conversion of PrP^C^ to PrP^Sc^ or act through established molecular targets

To define the mechanism of action of elacridar, we first examined its effects on PrP^C^ levels and localization. Treatment of uninfected N2a cells for 7 days with 1 µM elacridar, which is near the EC_50_ required to prevent prion propagation (Figure 1A; Table 1), had no significant effect on the level of PrP^C^ (Figure 2A, B). A modest, but significant, decrease in PrP^C^ levels was observed at higher concentrations, with reductions of 17 and 29% at 2.5 and 5 µM, respectively (Figure 2A, B). The subcellular distribution of PrP^C^ in these cells was unaffected by treatment with elacridar, as revealed by immunofluorescence staining (Figure 2C).

**Figure 2:**
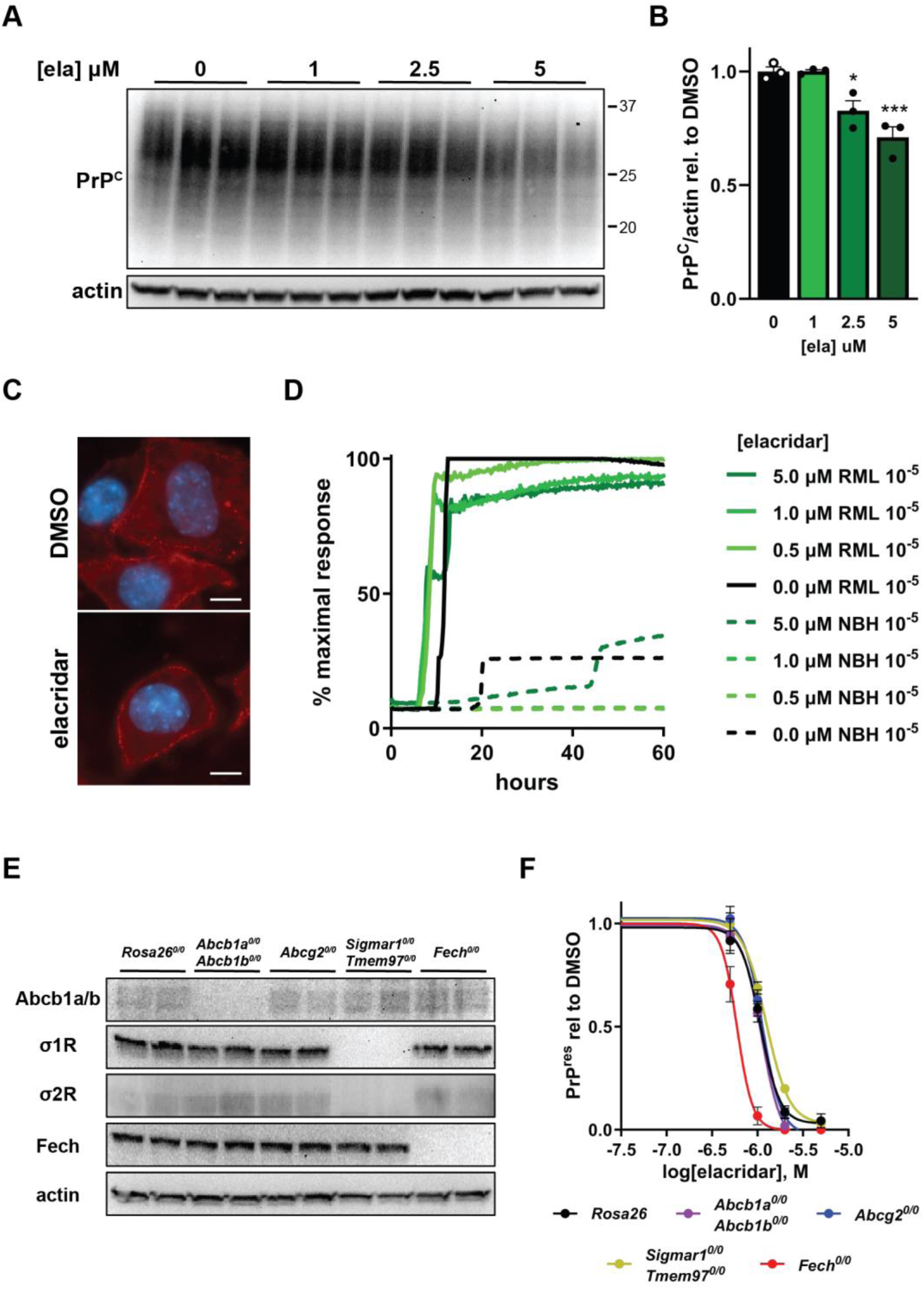
Elacridar does not alter PrP^C^, inhibit seeded amplification *in vitro*, or depend on previously identified molecular targets. **A)** Western blot of total levels of PrP^C^ following a 7-day exposure of uninfected N2a cells to elacridar. **B)** Levels of PrP^C^ ± SEM relative to DMSO treated cultures shown in **A** are plotted. p < 0.0001 = ****; p < 0.001 = ***; p < 0.01 = **; p < 0.05 = *; ns = not significant using one-way ANOVA and Dunnett’s multiple comparisons test. **C)** Cell surface levels of PrP^C^ are not altered by a 7-day exposure of uninfected N2a cells to 2.5 μM elacridar. Scale bar = 10 μm. **D)** Elacridar does not inhibit seeding activity of RML prions in a RT-QuIC assay. Uninfected (NBH) and prion infected (RML) brain homogenates were diluted to 10^−5^ for use as seeds. **E)** Western blot demonstrating successful knockout of *Abcb1a/Abcb1b*, *Sigmar1*, *Tmem97*, and *Fech*. **F)** Knockout cells were exposed to increasing concentrations of elacridar for 7 days, followed by western blotting for PK-resistant PrP. The ability of elacridar to reduce the levels of PrP^Sc^ was not affected by knockout of *Abcb1a/Abcb1b*, *Abcg2*, *Sigmar1* and *Tmem97*, or *Fech*.

To assess whether elacridar inhibits the conversion process through a direct interaction with PrP^C^ or PrP^Sc^, we utilized Real-Time Quaking-Induced Conversion (RT-QuIC), an *in vitro* seed amplification assay (31). We found that elacridar, at concentrations up to 5 µM, did not inhibit the seeding activity of RML prions in this assay (Figure 2D). It was not possible to assess higher concentrations of elacridar using RT-QuIC because of elacridar’s intrinsic fluorescence, with excitation/emission spectra overlapping with that of ThT (Supplementary Figure 3). However, at 5 µM, elacridar was in stoichiometric excess with respect to both the recombinant PrP^C^ substrate and PrP^Sc^ seed, making it unlikely that a specific effect would be observed at higher concentrations.

Elacridar was originally developed as an inhibitor of the MDR1 (*Abcb1a/Abcb1b* genes in mice; also known as P-glycoprotein) drug efflux pump, and has been found to interact with several other targets, including Abcg2 (another drug efflux pump known as BCRP1) (32), ferrochelatase (*Fech*, which catalyzes the last step in heme biosynthesis) (15), and the sigma receptors σ2R (*Tmem97*) (15,16) and σ1R (*Sigmar1*) (Supplementary Table 1). To investigate the role of these proteins in the anti-prion effects of elacridar, we performed CRISPR-Cas9 mediated gene knockout using ScN2a-RML cells. We performed a double knockout for *Abcb1a* and *Abcb1b*, and *Sigmar1* and *Tmem97*, and single knockout for *Abcg2* and *Fech*. Editing efficiency was assessed by DNA sequencing (Supplementary Figure 4A), and western blotting for the encoded proteins (Figure 2E). We were unable to detect Abcg2 by immunoblotting, and therefore relied on genomic analysis for confirmation of knockout (Supplementary Figure 4A). None of these gene knockouts altered the basal levels of PrP^Sc^ or the anti-prion effects of elacridar in prion-infected cultures, leading to the conclusion that these proteins do not mediate the anti-prion activity of elacridar (Figure 2F; Supplementary Figure 4B).

### Non-dividing cells are resistant to the anti-prion effects of elacridar

Cell division has been shown to significantly influence the anti-prion effects of some compounds. In particular, cell division typically enhances anti-prion potency (7–9), while inhibition of cell division has the opposite effect (9,33). This phenomenon has mechanistic implications regarding which stages of prion infection are targeted by a given compound. For example, the cellular pathways underlying prion propagation in dividing cells may be different from those responsible for maintaining infection in stationary cells. Moreover, cell division can influence the selection of drug-resistant prions *in vitro* (33).

We employed two strategies to examine the effects of elacridar in non-dividing cells. First, we utilized sodium butyrate (NaB) to inhibit division of ScN2a-RML cells. In dividing cells grown in the absence of NaB, elacridar at 2.5 µM began to lower the levels of PrP^Sc^ after 3 days of treatment. After 7 days, PrP^Sc^ was undetectable (Figure 3A, solid green line). In contrast, non-dividing cells grown in the presence of NaB were completely resistant to the anti-prion effects of elacridar (Figure 3A, dashed green line). This effect of NaB treatment on the action of elacridar was profound; even when challenged with concentrations as high as 10 μM, NaB treated cultures of ScN2a-RML cells showed no reduction in PrP^Sc^ levels (Figure 3B). Of note, these results were distinct from those reported in a similar experiment analyzing the effect of NaB on the ability of quinacrine, another anti-prion agent, to reduce PrP^Sc^ levels. In the presence of NaB, quinacrine caused a transient reduction in the amount of PrP^Sc^ at early time points, but PrP^Sc^ levels quickly rebounded as drug resistance emerged within the prion population (33).

**Figure 3:**
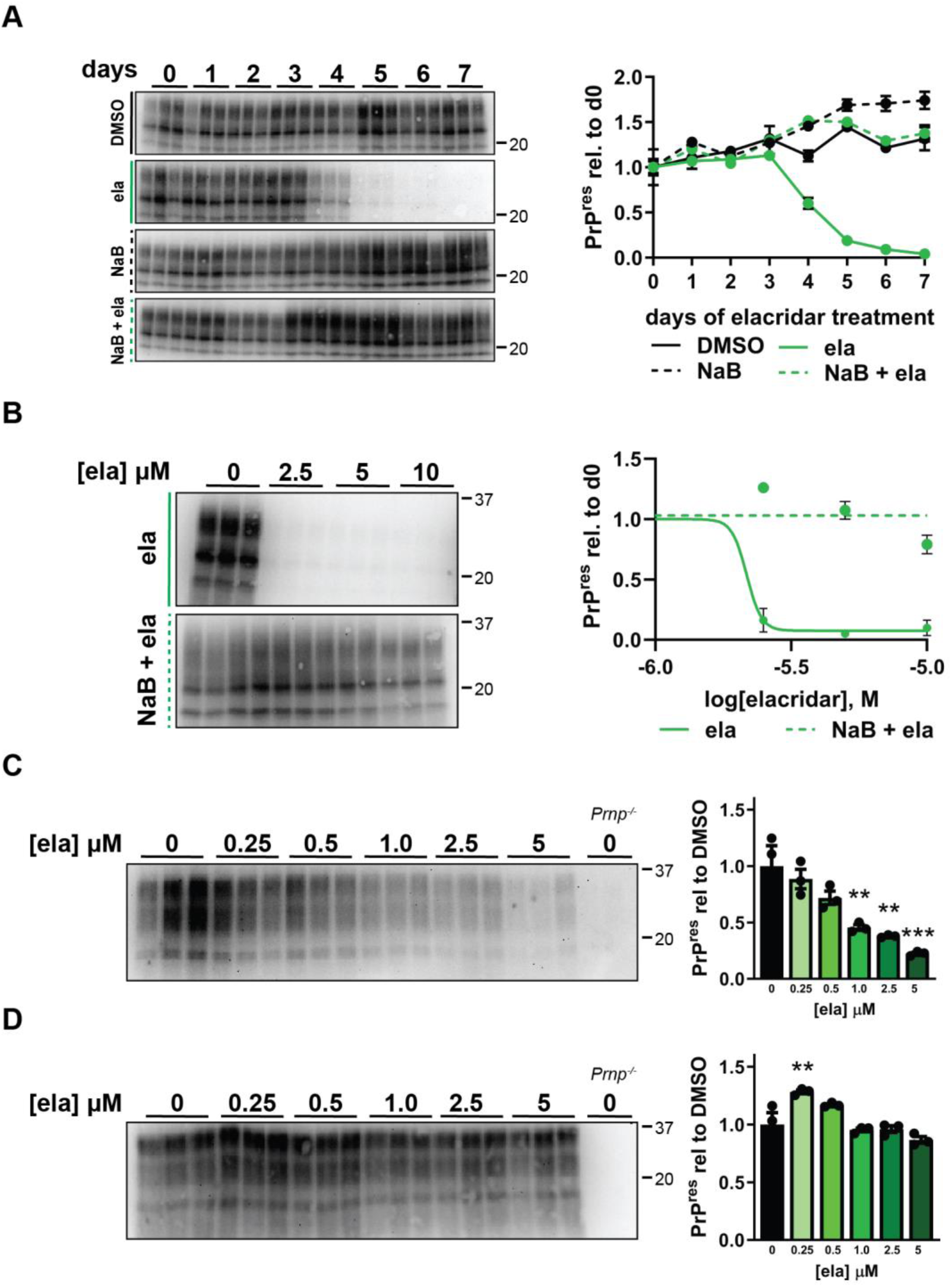
The anti-prion properties of elacridar are diminished in division-arrested cells. **A)** ScN2a-RML cells were treated for the indicated number of days with 2.5 μM elacridar, 10 mM sodium butyrate (NaB), or both and analyzed for PK-resistant PrP by western blotting. NaB treatment prevented the elacridar-induced reduction in PrP^Sc^ observed in cells treated with elacridar alone. **B)** ScN2a-RML cells were treated with increasing concentrations of elacridar with or without NaB and lysed after 7 days. NaB treatment prevented the elacridar-induced reduction of PK-resistant PrP at all concentrations tested. **C-D)** Cerebellar astrocytes infected with 22L were cultured in the presence of the indicated concentrations of elacridar for 5 days before western blot analysis for levels of PK-resistant PrP. Astrocytes were in either a proliferative state **(C)** or contact-inhibited state **(D)**. Elacridar reduces PrP^Sc^ in proliferative astrocytes with an EC_50_ = 647 nM (R^2^ = 0.8272), but has no effect on PrP^Sc^ in contact inhibited astrocytes. Level of PrP^Sc^ relative to untreated cultures ± SEM is plotted for both **C** and **D**. p < 0.0001 = ****; p < 0.001 = ***; p < 0.01 = **; p < 0.05 = *; ns = not significant using one-way ANOVA and Dunnett’s multiple comparisons test.

Because NaB is a non-selective agent that has a multitude of effects on cells, we analyzed the anti-prion properties of elacridar using a cell type that assumes a non-dividing state in the absence of pharmacological inhibition. For this purpose, we utilized primary cerebellar astrocytes infected with 22L prions *in vitro* (34). If allowed to grow to confluence, these cells cease proliferating due to contact-inhibition (35). Treatment with elacridar lowered the levels of PrP^Sc^ in sub-confluent, proliferating astrocyte cultures at concentrations similar to those found to be effective in proliferating ScN2a-RML cells (EC_50_ = 647 nM, Table 1, Figure 3C). However, as observed in NaB treated ScN2a-RML cells, the effectiveness of elacridar was dramatically diminished when applied to contact-inhibited cultures of prion-infected astrocytes (Figure 3D).

### Elacridar acts at an early stage of prion infection

The previous data demonstrates that elacridar has a greatly reduced ability to clear PrP^Sc^ from non-dividing cells with an established prion infection. We wondered whether elacridar might be more potent during the initial stages of prion exposure, before chronic infection is established. This hypothesis would be consistent with our observation that elacridar prevents retraction of dendritic spines on hippocampal neurons if applied within the first 3 hours after PrP^Sc^ exposure, but becomes progressively less effective at later times (Figure 1C-G).

First, we found that elacridar blocks the *de novo* infection of N2a cells by RML prions with an EC_50_ of 52.1 nM (R^2^ = 0.7665), more than 10x lower than the EC_50_ for the reduction of PrP^Sc^ in chronically infected N2a cells (Figure 4A; 1A; Supplementary Figure 5A; Table 1). Next, we tested the ability of elacridar to prevent the infection of post-mitotic cultures of C2C12 myotubes (36). To determine whether the efficacy of elacridar declined as the infection became established, we added elacridar at regular intervals following exposure to RML prions. When elacridar was present at the time of infection, or added during the subsequent three days, the levels of PrP^Sc^ at the end of the experiment were significantly reduced compared to untreated cultures (Figure 4B, Supplementary Figure 5B). In contrast, when elacridar addition was delayed by 4 or more days following the initial prion exposure, the final level of PrP^Sc^ was indistinguishable from that of untreated cultures, with the exception of a slight increase in cultures treated for 3 days (Figure 4B, Supplementary Figure 5B). These data indicate that elacridar is most effective during the early stages of prion infection, with diminished efficacy as the infection becomes established.

**Figure 4:**
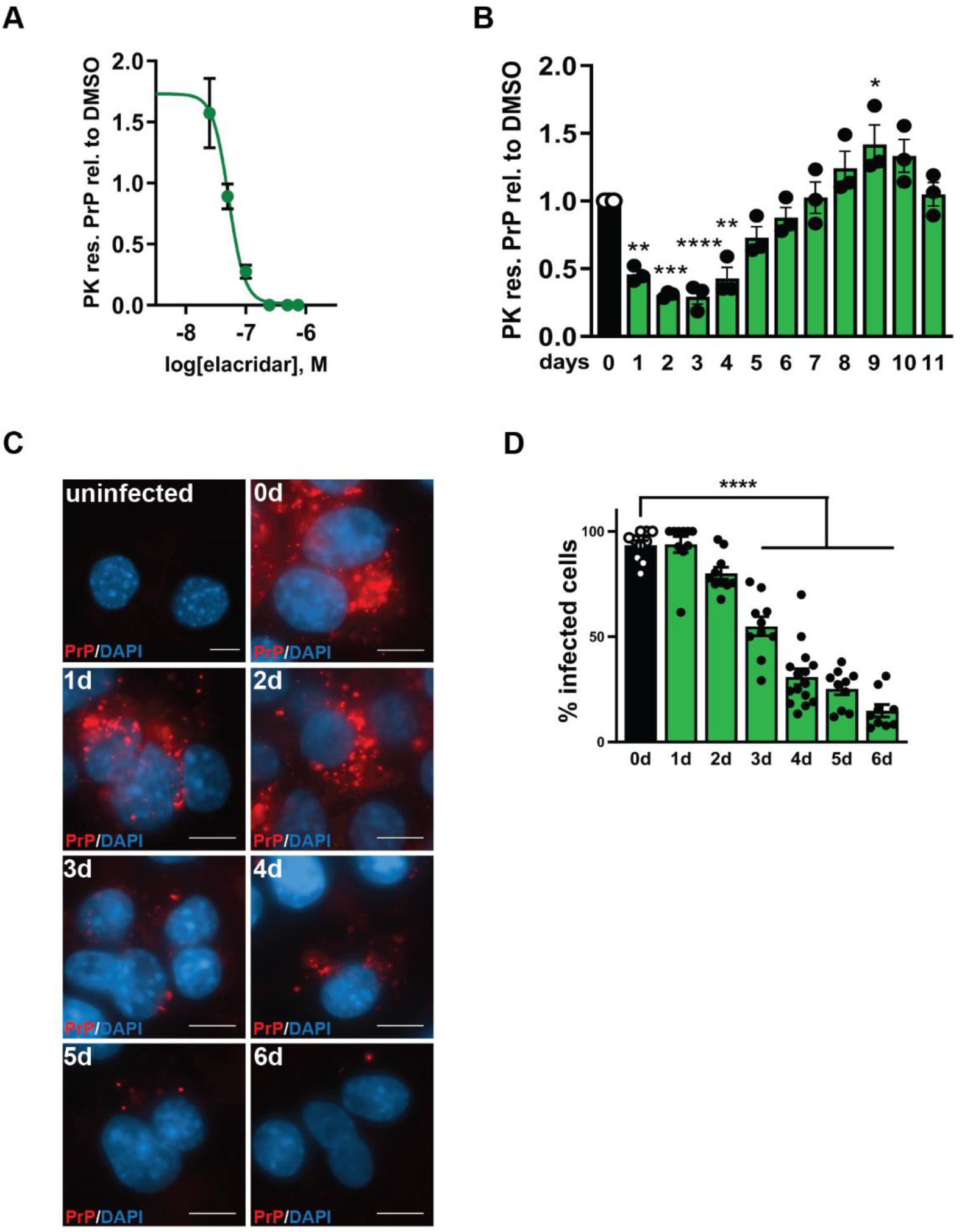
Elacridar prevents prion infection of naïve cells, and reduces cell-to-cell spread of prions. **A)** Dose response curve for the effect of elacridar on *de novo* prion infection. Elacridar was added at the time of initial infection of N2a cells by RML prions, and was present during the subsequent 4 passages, at which time cells were analyzed for PK-resistant PrP by western blotting. EC_50_ = 52.1 nM (R^2^ = 0.7665). **B)** Elacridar was ineffective at lowering the levels of PK-resistant PrP of non-proliferative C2C12 myotubes when added after the 4th day post infection with RML prions. Day of elacridar addition following infection is indicated. Levels of PK-resistant PrP relative to untreated cultures ± SEM is plotted. **C)** RML-ScN2a cells were incubated with 2.5 uM elacridar for the indicated number of days before exposure to PIPLC and GdnHCl to reveal PrP^Sc^ by immunostaining with D18. Uninfected N2a cells are shown as a negative control. **D)** Percentage of infected cells ± SEM shown in **C** is plotted. For all quantification, p < 0.0001 = ****; p < 0.001 = ***; p < 0.01 = **; p < 0.05 = *; ns = not significant using one-way ANOVA and Dunnett’s multiple comparisons test.

The spread of prion infection within a culture of dividing cells depends on horizontal transmission from infected to uninfected cells, as well as on vertical transmission from mother to daughter cells (37). If elacridar targeted the early steps of prion infection, we hypothesized that it would interfere with either one or both modes of prion spread within the culture as cells proliferated, thereby causing a decrease in the proportion of prion-infected cells over time. To test this prediction, we visualized PrP^Sc^ in ScN2a-RML cultures by immunofluorescence microscopy. We used phosphatidylinositol-specific phospholipase C (PIPLC) to release PrP^C^ and retain PrP^Sc^ (38), followed by guanidine denaturation to reveal antibody epitopes (39). The specificity of this approach for the detection of PrP^Sc^ is demonstrated by the absence of PrP staining of uninfected cells (Figure 4C; Supplementary Figure 6A). Over the course of elacridar treatment, we observed a continual decline in the percentage of infected cells, an effect that became significant after 3 days in culture (Figure 4C, D). Following six days of treatment, the number of infected cells was 14%, compared to 93% in DMSO-treated cultures (Figure 4C, D; Supplementary Figure 6B). This result suggests that elacridar prevents the infection of naïve cells that is required to maintain prion infection in culture.

In a final experiment, we directly analyzed the effect of elacridar on the initial conversion of PrP^C^ to PrP^Sc^ following *de novo* infection. To selectively monitor newly formed PrP^Sc^, and distinguish it from inoculum-derived PrP^Sc^, we utilized cells expressing PrP^C^ tagged with an epitope recognized by the 3F4 antibody (40). Following exposure of N2a cells to 22L prions, we found that 3F4 positive PrP^Sc^ appeared within one day, and persisted through day 5 (Figure 5A, B; black line). This result is consistent with previously published studies which have used this strategy to distinguish acute PrP conversion from chronic prion infection (41). The inclusion of elacridar in the inoculum, and in subsequent media changes, produced a very different result. Elacridar did not prevent the initial appearance of 3F4-positive PrP^Sc^ at day 1, but caused a gradual decrease in this species over the subsequent 4 days, such that it is undetectable by day 5 (Figure 5A, B; green line). In analogous experiments using RML-infected CAD5 cells expressing 3F4-tagged PrP^C^, elacridar completely prevented the formation of 3F4-positive PrP^Sc^ (Figure 5C, D), likely reflecting slower kinetics of PrP^Sc^ formation and/or turnover in CAD5 compared to N2a cells. These data clearly demonstrate that elacridar acts on an early step of prion infection, and suggests that it may influence the degradation of nascent PrP^Sc^ before it can establish a chronic infection.

**Figure 5:**
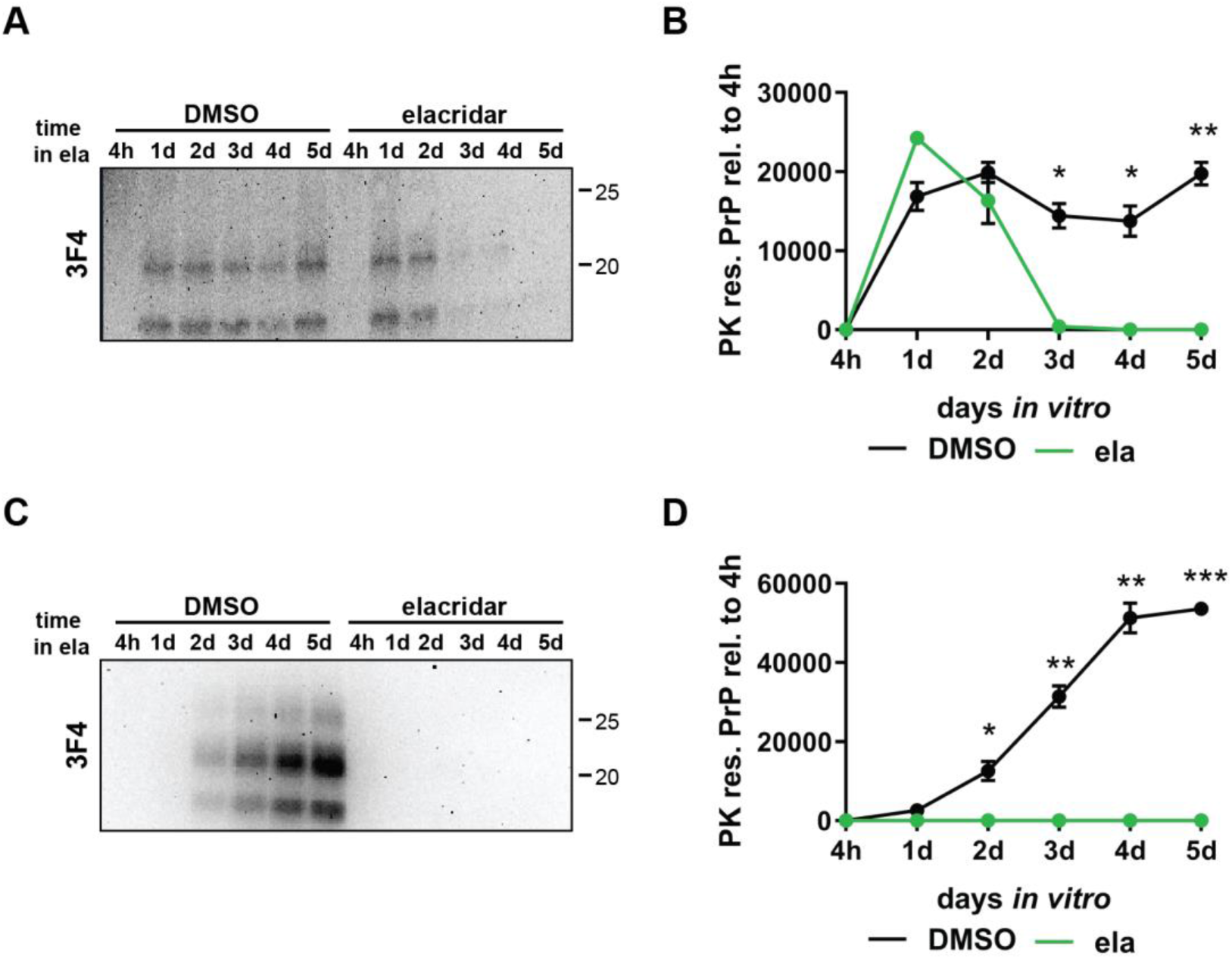
Elacridar prevents prion infection downstream of the initial PrP^C^ to PrP^Sc^ conversion event. N2a and CAD5 cells expressing mouse PrP with the 3F4 epitope were used to selectively monitor the conversion of endogenous PrP^C^ to PrP^Sc^ at early times following prion exposure. **A)** Western blot using the 3F4 antibody following exposure of N2a cells to 22L prions. **B)** Plot of 3F4 signal intensity in **A** with 4h set to 0. **C)** Western blot using the 3F4 antibody following exposure of CAD5 cells to RML prions. **D)** Plot of 3F4 signal intensity in **C** with 4h set to 0. For all quantifications, p < 0.0001 = ****; p < 0.001 = ***; p < 0.01 = **; p < 0.05 = *; ns = not significant using Student’s t-test with Welch’s correction.

Taken together, each of the experiments described in this section support the conclusion that elacridar interferes with an early event of prion processing by cells, prior to the establishment of a persistent infection.

### Elacridar accumulates in and activates lysosomes

The experiments shown in Figure 5 raised the possibility that elacridar is enhancing degradation of newly formed PrP^Sc^. To investigate further the cellular mechanisms underlying such an effect, we carried out transcriptomic analysis of N2a cells treated for 7 days with three different concentrations of elacridar (250, 700, and 2500 nM) which span the EC_50_ of elacridar’s inhibitory effect on PrP^Sc^ levels in proliferative cell lines (Figure 1A; Table 1). At all three concentrations, there was a significant enrichment for gene ontology (GO) terms related to lipid, sphingolipid, and sterol metabolic processes (Figure 6A-C; Supplementary Tables 2-4). This result was expected, given the known interaction of elacridar with Abcb1a/b, Abcg2, and the sigma receptors, which are involved in lipid transport and metabolism (42–44). Beginning at elacridar treatments of 700 nM, and increasing at 2500 nM, additional GO terms were returned related to regulation of cellular pH and lysosomal acidification, suggesting that lysosomes and other acidic organelles could play a role in the anti-prion activity of elacridar (Figure 6B, C; Supplementary Tables 3, 4). A heat map of the fold change in the expression of the leading edge genes driving the inclusion of GO terms related to cellular pH and lysosomal acidification demonstrates that their expression increases upon elacridar treatment in a dose dependent manner (Figure 6D). Because the protein products of many of these genes are found in the lysosome, including four subunits of the vacuolar H+-ATPase (*Atp6v0c*, *Atp6v0a1*, *Atp6vap2*, and *Atp6vap1*; Figure 6D, Supplementary Tables 3, 4), we hypothesized that elacridar may act through alteration of the function of this organelle, possibly though increasing lysosomal acidification. When we interrogated our transcriptomic data using a canonical set of 56 genes associated with lysosomal/autophagosomal structure or function (45), we found that elacridar at 700 nM or 2500 nM caused statistically significant increases in expression of genes encoding lysosomal hydrolases (e.g. *Naglu*, *Gba*, *Ctsa*, *Hexb*), lysosomal membrane proteins (e.g., *Ctns*, *Cln3*), and V-ATPase subunits (e.g., *Atp6v0c*, *Atp6v0b*, *Atp6v0a1*, *Atp6v0e*, *Atp6ap1*), consistent with expansion of the lysosomal compartment (Supplementary Table 5).

**Figure 6:**
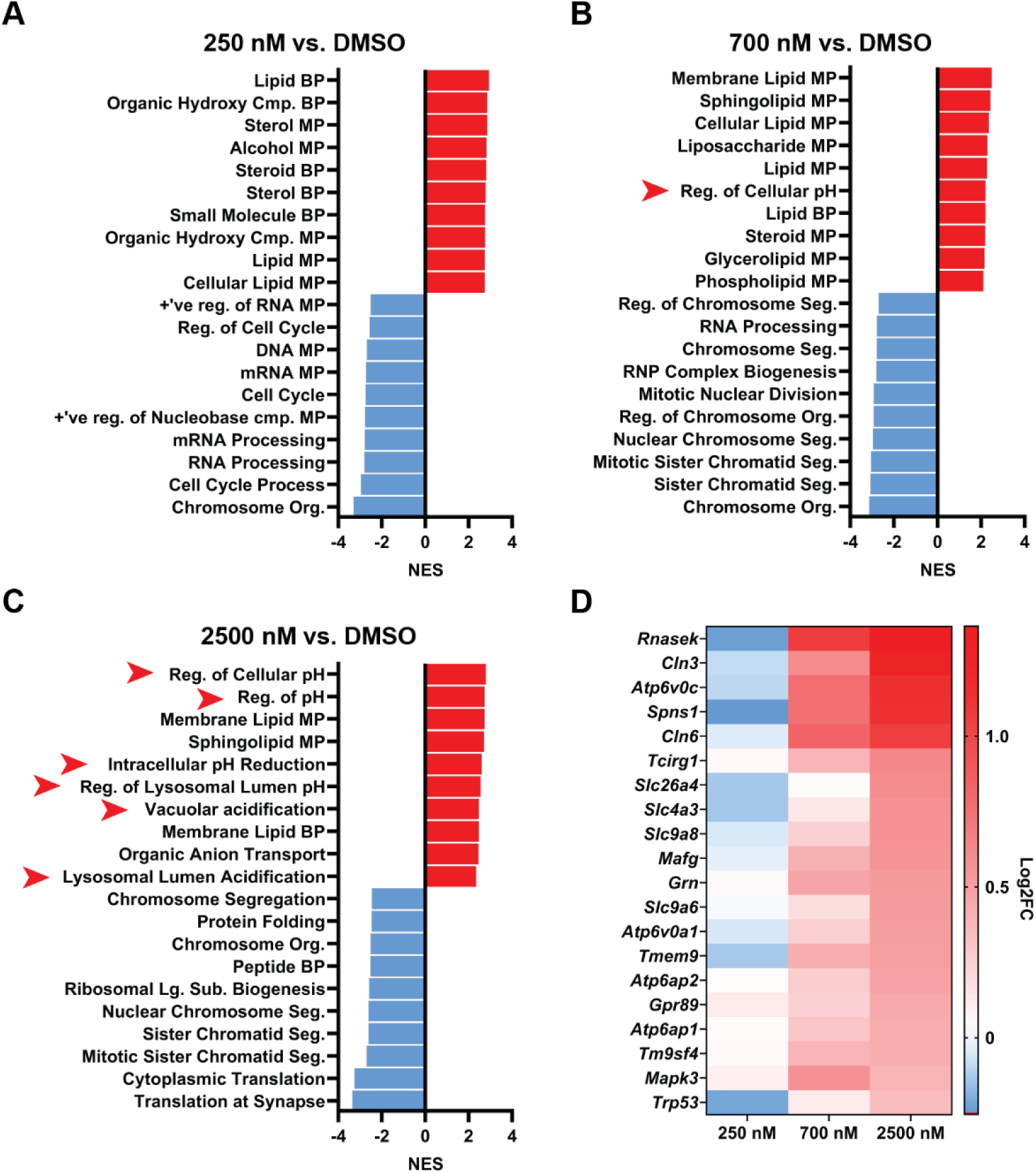
Elacridar upregulates lysosome-associated transcriptional programs. N2a cells were treated for 7 days with three different concentrations of elacridar before total RNA isolation and transcriptomic analysis by RNAseq. **A-C)** The top 10 Gene Ontology Normalized Enrichment Scores for biological processes with a padj value < 0.001 in each direction are shown following treatment with elacridar at **A)** 250 nM, **B)** 700 nM, and **C)** 2500 nM. Red arrowheads indicate terms related to the regulation of cellular pH. **D)** Heatmap of leading edge genes driving the inclusion of cellular pH GO terms.

To investigate the effects of elacridar on lysosomes at a cellular level, we took advantage of the intrinsic fluorescence of elacridar (ex: 450 nm, em: 480 nm; Supplementary Figure 3) to visualize its subcellular localization. Following application to N2a cells, we observed intracellular accumulations of elacridar that colocalized with LysoTracker Red DND-99 (LTR99), demonstrating that the compound concentrates in acidic intracellular compartments (Figure 7A, B). We also observed colocalizaton of elacridar with DQ-Red-BSA, the fluorescence of which is self-quenching until the BSA is degraded, thereby demonstrating that the compartments in which elacridar is accumulating are proteolytically active (Figure 7C, D). Bafilomycin A1, which causes an increase in lysosomal pH though inhibition of the vacuolar H+-ATPase (46), prevented the intracellular accumulation of elacridar and LTR99, and also diminished proteolysis of DQ-Red-BSA, further supporting the conclusion that elacridar localizes to lysosomes (Figure 7A iv, viii; C iv, viii). We noted that elacridar increased the intensity of LTR99 staining and the number of DQ-Red-BSA puncta/cell (Figure 7), consistent with enhancement of lysosomal function. The targeting of elacridar to the lysosome was not due to interaction with its known targets, as demonstrated by elacridar accumulation in *Abcb1a* + *Abcb1b*, *Abcg2*, *Sigmar1 + Tmem97*, and *Fech* knockout cell lines (Supplementary Figure 7). To test the effect of elacridar on autophagy, we measured its effect on the proportion of lipidated LC3, which is associated with autophagosomes. We found that elacridar significantly increased the ratio of lipidated to unlipidated LC3, suggesting enhancement of autophagosome formation (Supplementary Figure 8).

**Figure 7:**
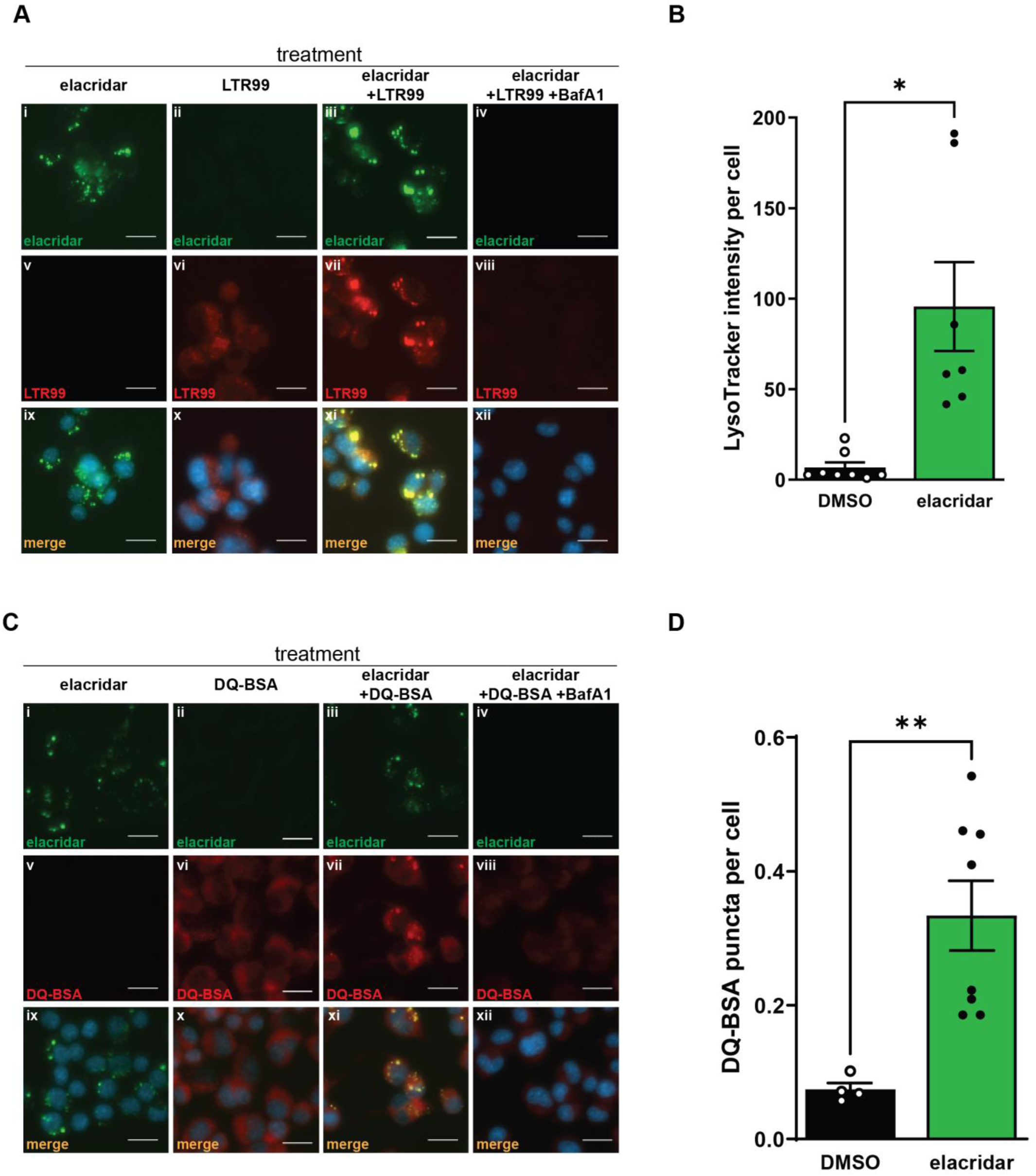
Elacridar accumulates in lysosomes and stimulates their proteolytic activity. **A)** Elacridar accumulates in intracellular compartments that co-stain with LysoTracker Red 99 (LTR99), and this is prevented by the co-administration of bafilomycin A1 (BafA1). N2a cells were incubated with elacridar for 3 days prior to imaging. **B)** Average LTR99 intensity per cell ± SEM shown in **A vi** (DMSO) and **A vii** (elacridar) is plotted. **C)** Elacridar co-localizes DQ-BSA fluorescence, which is prevented by the co-administration of BafA1. N2a cells were incubated with elacridar for 3 days prior to imaging. **D)** Average DQ-BSA intensity per cell ± SEM **C vi** (DMSO) and **C vii** (elacridar) is plotted. Scale bars = 20 µm. For all quantification, p < 0.0001 = ****; p < 0.001 = ***; p < 0.01 = **; p < 0.05 = *; ns = not significant using Student’s t-test with Welch’s correction.

The lysosomal accumulation of elacridar was also prevented by the application of ammonium chloride, a weak base that directly neutralizes lysosomal pH (Supplementary Figure 9). This result is consistent with chemical nature of elacridar as a cationic amphiphilic drug, or CAD. CADs are known to accumulate in acidic intracellular compartments in an uncharged state, and then become trapped there following protonation and interaction with membranes (47). We compared the localization of elacridar with quinacrine, another CAD that has anti-prion properties and is intrinsically fluorescent (48,49). We observed that the intracellular distribution of quinacrine was distinct from that of elacridar, accumulating in in a more diffuse pattern, and having minimal impact on LTR99 and DQ-Red-BSA fluorescence (Supplementary Figure 10). These results suggest that the anti-prion mechanisms of elacridar and quinacrine are distinct.

Because they accumulate in endolysosomal compartments, CADs have been associated with disruptions in lipid processing, resulting in a phenomenon known as drug induced phospholipidosis (DIPL) (50). While elacridar has not been associated with DIPL (51), this has not been investigated using N2a cells. We therefore compared elacridar and quinacrine to amiodarone, a third anti-prion compound and potent inducer of phospholipidosis (14,51,52). Based on LipidTox Red staining, a commonly used readout of DIPL (51), we found that that elacridar treatment of N2a cells does not result in DIPL, in contrast to quinacrine which strongly induced phospholipidosis at levels comparable to amiodarone (Figure 8A, B). This result is consistent with the conclusion that elacridar and quinacrine exert their anti-prion activity via different mechanisms.

**Figure 8:**
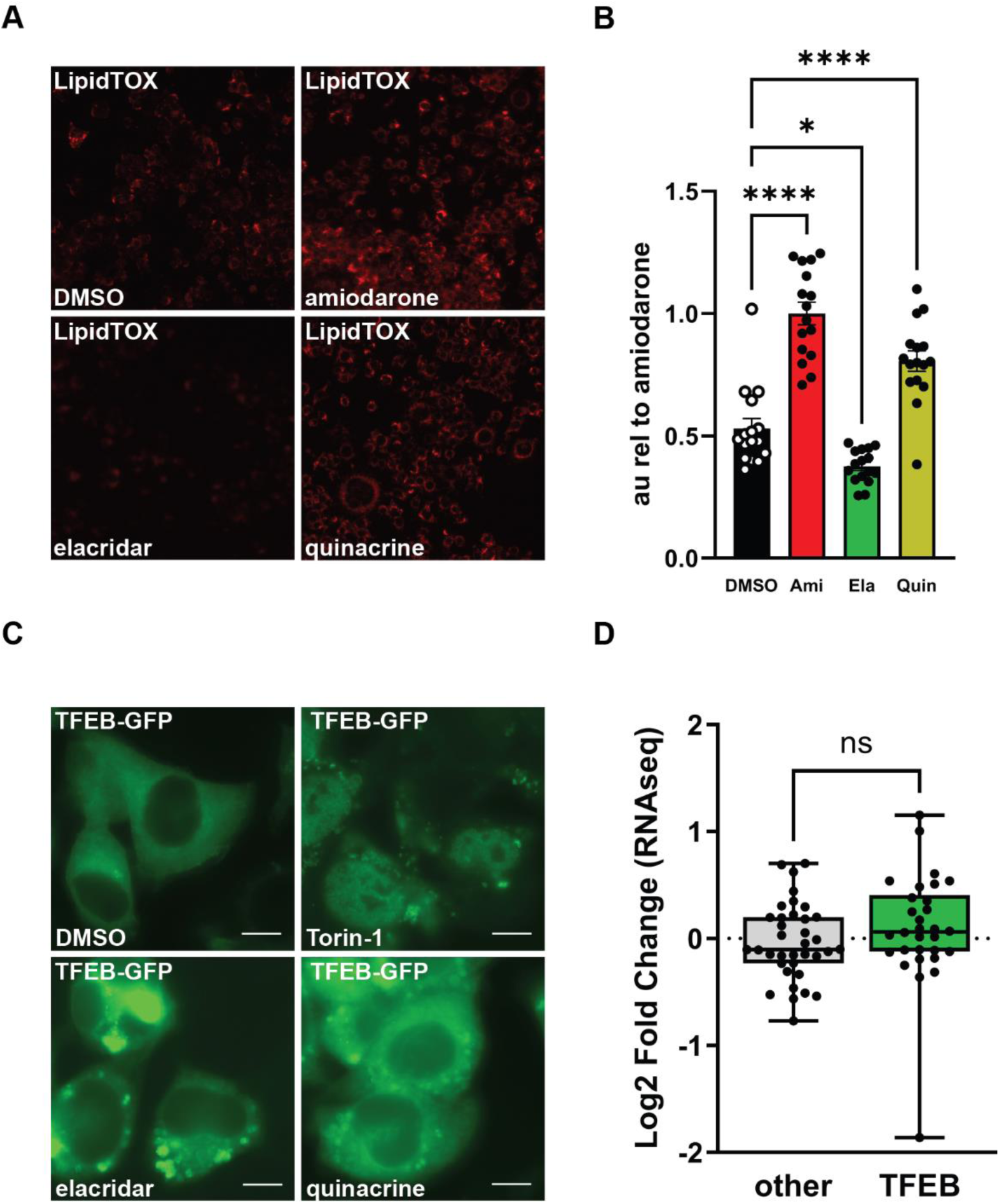
Elacridar does not induce phospholipidosis or cause nuclear translocation of TFEB. **A)** LipidTOX red staining reveals the induction of phospholipidosis by amiodarone and quinacrine, but not elacridar. N2a cells were incubated with the compounds for 24 hours prior to imaging. **B)** Intensity of LipidTOX fluorescence relative to amiodarone treated cultures ± SEM shown in **A** is plotted and statistical analysis performed using one-way ANOVA and Dunnett’s multiple comparisons test. **C)** TFEB-GFP localizes to the nucleus of N2a cells when cultured in the presence of Torin-1 but not elacridar or quinacrine. N2a cells transfected to express TFEB-GFP were incubated with the compounds for 24 hours prior to imaging. Scale bar = 10 µm. **D)** Log2 fold change of TFEB responsive genes (TFEB) vs non-TFEB responsive genes (other) taken from RNA-seq analysis of N2a cells treated with 2500 nM elacridar. Lists of genes were taken from (56). Average log2 fold change ± SEM is plotted and statistical analysis performed using Student’s t-test with Welch’s correction. For all quantification, p < 0.0001 = ****; p < 0.001 = ***; p < 0.01 = **; p < 0.05 = *; ns = not significant.

The lysosomal dysfunction induced by some CADs can lead to a compensatory activation of transcription factor EB (TFEB) which, upon dephosphorylation, translocates to the nucleus to coordinate the expression of an array of genes required for lysosomal biogenesis (53,54). We used a GFP-tagged version of TFEB to monitor this translocation. In a positive control experiment, we observed that Torin-1, an mTOR inhibitor, caused dramatic translocation of TFEB-GFP into the nucleus of N2a cells (55). In contrast, no change in TFEB-GFP translocation was detected after application of elacridar or quinacrine (Figure 8C). Further, we did not observe an enrichment of transcriptomic signatures associated with TFEB activation in our RNA-seq dataset (56,57) (Figure 8D). We, therefore, conclude that elacridar’s effect upon lysosomes occurs via a TFEB-independent pathway.

### Elacridar prevents the aggregation of α-synuclein and tau

Because of its activating effects on lysosomes, we hypothesized that elacridar would also prevent the propagation of protein aggregates associated with other protein misfolding neurodegenerative diseases. To test this, we measured the effect of elacridar on the seeding activity of misfolded α-synuclein and tau prions using HEK293T biosensor cells expressing YFP tethered to either full-length α-synuclein with the A53T mutation (α-syn140*A53T–YFP) (58,59) or the repeat domain of tau with the P301L and V337M mutations (TauRD(LM)– YFP) (60,61). α-syn140*A53T–YFP cells were plated in the presence of elacridar or DMSO before exposure to PTA precipitates from the brains of TgM83^+/−^ mice, which express A53T human α-synuclein (62), that had been inoculated with control brain homogenate (TgM83^+/−^ (−)), or brain homogenate from a patient with multiple system atrophy (MSA) (TgM83^+/−^ (+)) (63). TauRD(LM)–YFP cells were similarly challenged with PTA precipitates from the brains of aged C57BL/6J mice, or symptomatic Tg2541^+/+^ mice, which express the P301S mutation of tau associated with frontotemporal dementia (64). As expected, biosensor cells exposed to PTA precipitates of TgM83^+/−^ (+) or Tg2541^+/+^ brain homogenates developed intracellular aggregates within three days (38.3 ± 2.8%, α-syn; 32.9 ± 2.7%, tau) that were absent in cells challenged with precipitates from control brains (1.6 ± 0.4%, α-syn; 3.2 ± 0.4%, tau) (red arrows, Figure 9A i-ii, B, C i-ii, D). Despite the strong fluorescent signal from YFP, it was possible to identify intracellular accumulations of elacridar, which likely contribute to slightly increased aggregate counts in elacridar-treated cells that were not exposed to α-synuclein or tau prions (5.6 ± 0.9%, α-syn; 8.2 ± 0.8%, tau) (white arrows, Figure 9A iii, B, C iii, D). However, when elacridar treated cells were challenged with precipitates from TgM83^+/−^ (+) or Tg2541^+/+^ brains, there was a clear reduction in the aggregation of the α-synuclein and tau YFP biosensors compared to DMSO treated cells (11 ± 1.2%, α-syn; 10 ± 1.0%, tau) (Figure 9A iv, B, C iv, D). Thus, in addition to its effects on PrP^Sc^, elacridar is able to prevent seeded aggregation of α-synuclein and tau.

**Figure 9:**
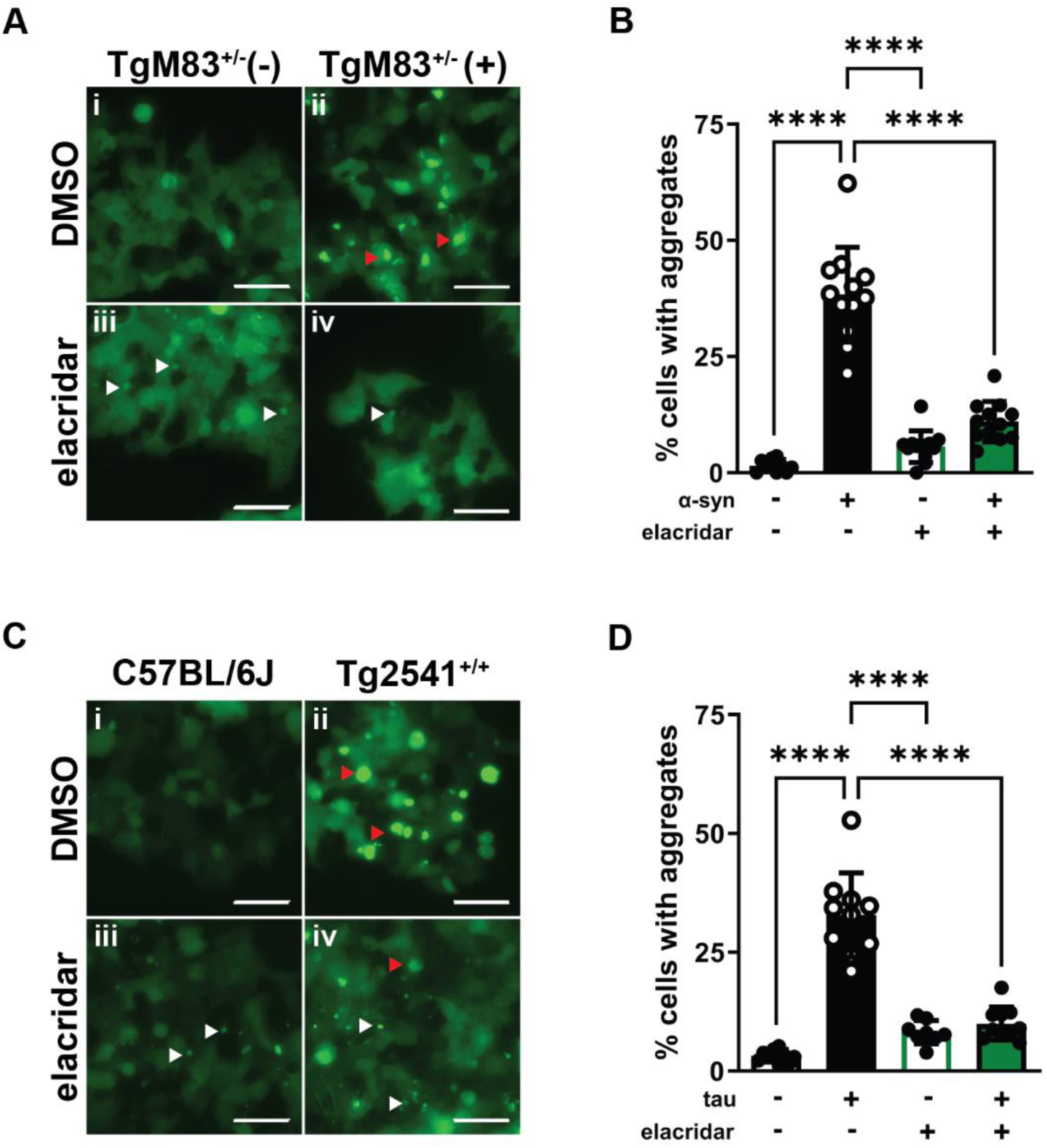
Elacridar inhibits intracellular seeding of α-synuclein and tau aggregates. **A)** HEK293T cells expressing α-syn140*A53T–YFP were exposed to PTA precipitates from TgM83^+/−^ mice inoculated with control brain (−) or brain homogenate from a patient with multiple systems atrophy (MSA, +) in the presence or absence of 5 µM elacridar. **B)** Percentage of cells with aggregates ± SEM shown in **A** is plotted. **C)** HEK293T cells expressing TauRD(LM)–YFP were exposed to PTA precipitates from aged C57BL/6J mice, or symptomatic Tg2541^+/+^ mice in the presence or absence of 5 µM elacridar. **D)** Percentage of cells with aggregates ± SEM shown in **C** is plotted. YFP shown in green, scale bars = 50 µm. Red arrows indicate misfolded α-syn and tau, white arrows indicate fluorescent signal from elacridar. For all quantification, p < 0.0001 = ****; p < 0.001 = ***; p < 0.01 = **; p < 0.05 = *; ns = not significant using one-way ANOVA and Dunnett’s multiple comparisons test.

## Discussion

In the present work, we have undertaken a detailed mechanistic investigation of the anti-prion activity of elacridar, a compound that has previously been used clinically as an adjuvant in cancer chemotherapy. We discovered that elacridar prevents prion propagation and synaptotoxicity at submicromolar concentrations *in vitro* and determined that this activity is unrelated to changes in the levels or subcellular localization of PrP^C^, direct interference in the conversion of PrP^C^ to PrP^Sc^, or interaction with its previously characterized molecular targets. We found that elacridar acts by stimulating the activity of lysosomes, potentially enhancing PrP^Sc^ degradation, and that it is most effective early in the infectious process, downstream of the initial PrP conversion event. Finally, we have shown that the inhibitory effects of elacridar extend to prions of α-synuclein and tau.

### Elacridar boosts lysosomal activity

Several lines of evidence demonstrate that elacridar enhances lysosomal activity. The drug accumulates in lysosomes by virtue of its cationic amphipathic chemical properties, colocalizing with and increasing the intensity of fluorescent accumulations of LTR99 and DQ-Red-BSA. Consistent with expansion and enhanced functional activity of the lysosomal compartment, elacridar activates transcription of genes encoding lysosomal hydrolases, membrane proteins, and subunits of the V-ATPase, and upregulates pathways related to lipid and sterol metabolism, and lysosomal pH. Upregulation of V-ATPase subunits is particularly prominent, raising the possibility that a major mechanism of elacridar’s action may be enhanced acidification of lysosomes. This effect sets elacridar apart from other CADs that primarily impact cholesterol and lipid homeostasis without directly altering the regulation of lysosomal pH (65). Elacridar also increases the ratio of lipidated/unlipidated LC3, suggesting an enhancement of autophagosome formation, although additional studies will be required to fully document how the elacridar alters autophagic pathways.

Interestingly, the effects of elacridar on lysosomes occur without nuclear translocation of TFEB, the “master transcriptional regulator” of lysosomal biogenesis (54,66–69). Translocation of unphosphorylated TFEB into the nucleus is generally required for upregulation of the Coordinated Lysosomal Expression and Regulation (CLEAR) gene network (45). There are other examples of TFEB-independent regulation of CLEAR genes (70,71), but further work will be necessary to fully elucidate the role of the CLEAR gene network in the action of elacridar.

### Elacridar enhances clearance of PrP^Sc^

Presumably as a consequence of increased lysosomal and autophagic activity, elacridar enhances clearance of PrP^Sc^. The most direct evidence for this conclusion comes from experiments designed to selectively monitor the production of nascent PrP^Sc^ after infection of N2a cells. In those experiments, elacridar did not prevent the initial appearance of PrP^Sc^, but prevented its accumulation. Our data do not rule out a direct inhibitory effect on the conversion of PrP^C^ to PrP^Sc^ in cells, but our *in vitro* experiments using the RT-QuIC seeding assay argue against this possibility.

Our results are consistent with previous studies showing that PrP^Sc^ is degraded primarily in lysosomal and autophagosomal pathways (72), and that activation of these pathways can reduce PrP^Sc^ levels (73). Several anti-prion compounds including imatinib (74), trehalose and rapamycin (75), metformin (76), and FK506 (77), are reported to enhance degradation of PrP^Sc^ via autophagy (78). Whether all of these compounds affect PrP^Sc^ metabolism via similar mechanisms is unclear. The accumulation of elacridar in lysosomes suggests a direct effect on this organelle, potentially setting it apart from other compounds that enhance PrP^Sc^ degradation by influencing upstream pathways.

### Elacridar acts early in prion infection

We have shown here that elacridar is most potent when applied to cells at the time of initial infection, and is ineffective against established prion infection in non-dividing cells, such as C2C12 myotubes, stationary phase astrocytes, and N2a cells treated with a mitotic inhibitor. To explain these results, we hypothesize that PrP^Sc^ accumulation in lysosomes progressively impairs their function, so that compounds like elacridar, which boost lysosomal function, become less effective as the infection proceeds. This idea is consistent with studies showing that accumulation of Aβ and α-synuclein aggregates in lysosomes and autophagosomes damages their integrity, leading to exposure of their contents to the cytoplasm, and eventually resulting in cell death (79,80). In rapidly dividing cells, prion persistence requires cell-to-cell spread through either horizontal or vertical transmission, providing an opportunity for elacridar to act.

### The anti-prion mechanism of elacridar is distinct from that of other lysosomotropic amines

Lysosomotropic agents, including quinacrine, have long been known to possess anti-prion properties (48). Elacridar and quinacrine share chemical properties that define them as CADs, but there are important mechanistic distinctions between the anti-prion effects of these molecules that are borne out by our investigations. Most importantly, quinacrine, unlike elacridar, is a strong inducer of phospholipidosis, a phenomenon characterized by a reversible accumulation of phospholipids due to impaired lysosomal lipid catabolism and cholesterol export (81). The anti-prion effects of quinacrine have been associated with the redistribution of cholesterol in cultured cells, which could be related to its ability to induce DIPL (52). This mechanism may also apply to other lysosomotropic, anti-prion compounds (48) that have been reported to induce phospholipidosis, including tilorone (82), chloroquine (83), and suramin (84). Elacridar and quinacrine also display differences in their cellular localization based on their intrinsic fluorescence, suggesting non-identical sites of action.

### What is the relevant molecular target of elacridar?

Several molecular targets of elacridar have been previously documented. The drug was originally identified as a non-competitive inhibitor of ABCB1 (also known as MDR1 and P-glycoprotein), a transporter that pumps xenobiotics out of cells. Based on this activity, elacridar has been used clinically as an adjuvant in cancer treatment to enhance cellular retention of chemotherapeutic agents (17). Other studies have demonstrated that elacridar binds to Tmem97 (σ2 receptor), Abcg2 (BCRP1), and ferrochelatase (15,32). We have shown that CRISPR editing to eliminate the expression of these proteins has no effect on the anti-prion effect of elacridar, definitively ruling them out as the molecular targets for this activity. Identification of the relevant target will require other approaches, such as an unbiased profiling with affinity-tagged versions of elacridar.

### Elacridar as a therapeutic for multiple prion diseases

A major finding of this work is that elacridar is capable of preventing the aggregation of tau and α-synuclein in biosensor cell lines exposed to brain-derived seeds. This result suggests that the expanded lysosomal capacity produced by elacridar enhances degradation of other self-propagating pathological protein assemblies besides PrP^Sc^. Lysosomes are important sites for the degradation of misfolded proteins that have been implicated in the pathogenesis of several neurodegenerative diseases, including Alzheimer’s disease (85), Parkinson’s disease (86), and others (87,88). Underscoring the importance of this organelle to the disease processes, lysosomal dysfunction itself is known to cause neurodegeneration (88), and genes associated with lysosomal storage disorders harbor mutations that are established risk factors for other neurodegenerative diseases (89–91). Activation or restoration of lysosomal function using small molecules has recently gained attention as a novel therapeutic modality for several neurodegenerative diseases (56,92–94), and the present work adds elacridar to the small but growing list of compounds with this activity.

Could elacridar also serve as an effective therapeutic for neurodegenerative diseases in clinical settings? Our observation that elacridar acts primarily during the initial steps of *de novo* prion infection, and is minimally effective on non-dividing cells with established infection, suggests that elacridar would be most useful as a preventative in cases of neurodegeneration caused by mutations in PrP, α-synuclein, and tau. In this scenario, elacridar could be administered to at-risk individuals in the pre-symptomatic phase, before prions have already accumulated to high levels. As a next step, it will be of interest to test elacridar in iPSC (95–97) and mouse models that express mutations in PrP, tau, and α-synuclein (62,64,98–100). Elacridar will be useful as a tool to dissect the mechanisms by which PrP^Sc^ and other prions are degraded by lysosomal and autophagic pathways, and may facilitate discovery of additional therapeutic compounds.

## Experimental Procedures

### Cell culture

Except where indicated, cells were grown in Opti-MEM (Gibco) supplemented with 10% FBS (Gemcell) and 100 U/ml each of penicillin/streptomycin (Gibco) in a humidified atmosphere of 5% CO_2_ at 37 °C. Chronic prion infection was achieved by exposure of confluent cultures to infected brain homogenate (0.1% final concentration) for 24 h before removal and subsequent passage. C2C12 myoblasts were cultured in DMEM (Gibco) containing 25 mM glucose and 1 mM pyruvate supplemented with 10% FBS and 100 U/ml each of penicillin/streptomycin. Once confluent, this media was switched to one containing 10% horse serum (replacing 10% FBS). After four days, horse serum was lowered to 1% and the cultures were exposed to 0.1% brain homogenate with daily media changes. For experiments with TFEB-GFP, cells were transfected with a pcDNA3.1 (+) vector using lipofectamine 3000 (Invitrogen) and selected using neomycin.

### Elacridar treatment of chronically infected cell lines

Cells were cultured with the indicated concentration of elacridar (Tocris) for a total of seven days, with passaging on the third day in a constant DMSO concentration of 0.1%. Cells were washed with cold PBS before lysis with cold cell lysis buffer (10 mM Tris pH 7.8, 100 mM NaCl, 0.5% NP-40, 0.5% Na deoxycholate and 0.1% SDS). MTT (Sigma) assays were performed in parallel, where indicated. Briefly, cells were incubated with 0.5 mg/ml MTT in PBS for 30 min at 37 ℃. Following aspiration of the MTT solution, formazan was solubilized with DMSO at 37 ℃ for 10 min before obtaining absorption readings at 570 nm using a Synergy H1 plate reader (BioTek). Signal was normalized to DMSO treatment and curves were fit by least squares regression using GraphPad software. Reagents were purchased from the following suppliers: Sodium butyrate and ammonium chloride from Sigma, LipidTOX, LysoTracker-Red-99, DQ-Red-BSA and PIPLC from Life Technologies, quinacrine, amiodarone, and torin-1 from Tocris.

### Proteinase K treatment

100 μg of protein, as determined by BCA assay (Pierce) was exposed to 10 μg/ml proteinase K (Roche) in a final volume of 250 μL in lysis buffer at 37 ℃ for 1 hour with 750 rpm shaking. 30 µL of 10x protease inhibitor (Pierce) was added and samples were centrifuged at 20817 x g for 1 hour at 4 ℃. Supernatant was removed and pellets were resuspended in 1x Laemmli buffer (BioRad).

### Western blot

Protein samples were incubated at 100 ℃ for 10 minutes in 1x Laemmli buffer (BioRad) and loaded into 12% Criterion TGX Precast Protein Gels and run at 200V for 42 min. Proteins were transferred to PVDF membranes for 45 min at 115V before washing in 0.1% TBST and blocking in 5% non-fat milk in 0.1% TBST for 1 hour. Primary antibodies used: D18 (anti-PrP(101), prepared in-house), 3F4 (anti-PrP, Millipore), σ1R (B-5, Santa Cruz Biotechnology), σ2R (26444-1-AP, Proteintech), Abcb1a/b (MA1-26528, ThermoFisher), Fech (A-3, Santa Cruz Biotechnology), LC3 (M186, MBL), β-actin (AC-74, Millipore Sigma). All HRP-conjugated secondary antibodies were purchased from BioRad. Membranes were visualized using ECL (Millipore). Quantification was performed using ImageJ and plotted using GraphPad software.

### Prion purification

Prion enrichment was performed as described (28). Briefly, 200 μl of 10% (w/v) RML infected brain homogenate was incubated with 100 μg/ml (final concentration) of pronase E (Sigma) for 30 min at 37 °C at 800 rpm. 10 mM (pH 8) EDTA, 2% sarkosyl in D-PBS, and 50 U/ml Benzonase (final concentrations) was added and incubated for 10 min at 37 °C. Samples were brought to 0.3% (w/v) NaPTA (pH 7.4) and incubated at 37 °C for 30 min. Samples were then mixed with iodixanol to a final concentration of 35% (w/v) while keeping NaPTA at 0.3% (w/v). Samples were centrifuged at 16, 100 x g for 90 min. The upper layer was recovered and filtered with a 0.45 μm pore size Durapore membrane Ultrafree-HV microcentrifuge filtration unit (Millipore). Samples were mixed with an equal volume of 0.3% NaPTA and 2% sarkosyl, incubated at 37 °C for 10 min, and centrifugation for 90 min at 16, 100 x g. The pellet was resuspended in D-PBS with 17.5% iodixanol and 0.1% sarkosyl. Samples were washed twice and resuspended in D-PBS with 0.1% (w/v) sarkosyl, pooled and stored as aliquots at −80 °C. Samples were analyzed by silver stain (per manufactures instruction; Pierce) and immunoblotting with and without PK digestion.

### Scrapie cell assay

RK13 cells expressing murine PrP infected with RML prions (RKM7+) and RK13 cells expressing ovine (VRQ) PrP infected with SSBP/1 prions (RkOv+) were cultured with elacridar for 7 days in DMEM supplemented with 10% FBS and 100 U/ml each of penicillin/streptomycin. 20,000 cells were plated onto activated ELISpot plates and exposed to 5 µg/ml PK in 50 mM Tris-HCl, pH 8.0, 150 mM NaCl, 0.5% sodium deoxycholate, and 0.5% Igepal CA-630 for 90 min at 37 °C before inactivation with 2 mM PMSF. Following PBS washes, membranes were blocked with SuperBlock Blocking Buffer (Thermo) and incubated with D18 antibody overnight at 4 ℃ before incubation with an alkaline phosphatase–conjugated goat anti-human IgG (Southern Biotech). Spots were developed with BCIP/NBT substrate (Roche) before imaging and quantification using a CTL Immunospot Imager.

### Dendritic spine retraction assay

Cultures of hippocampal neurons were prepared as described previously (24,26). Neurons were exposed to compounds or vehicle for 2 h before the application of 4.4 µg/ml of enriched prion preparations (or a mock-purified control sample from non-infected brains) for 24 h. 4% paraformaldehyde was used for fixation and neurons were stained with Alexa Fluor 488-phalloidin (Thermo Fisher) to visualize actin in dendritic spines. Images were acquired using a Zeiss 700 confocal microscope with a 63x objective using Zen software. Number of dendritic spines was determined using ImageJ software. Spine number was normalized to the measured length of the dendritic segment to give the number of spines/μm.

### Fluorescence microscopy

To reveal PrP^Sc^, cells were plated on coverslips and treated with elacridar for the indicated time period and exposed to PIPLC for 4 hours before processing as described (102). Cells were washed with PBS and fixed using 4% paraformaldehyde in PBS pH 7.4 for 12 min at room temperature. Cells were then washed with PBS before denaturation with 3M GdnHCl for 10 min. Following extensive washing with PBS, cells were then incubated in 10 ug/ml D18 for 1 hour at room temperature in a 1x solution of SuperBlock Blocking Buffer (Thermo). Cells were then washed 3x with PBS before incubation with Alexa Fluor-594 goat anti human secondary (Invitrogen) in the presence of 1 ug/ml Hoechest and 1x SuperBlock Blocking Buffer. Cells were washed 3x with PBS and mounted. In other experiments, LysoTracker Red DND-99 (Thermo) and DQ-Red-BSA (Thermo) were added to cell media at a final concentration of 50 nM and 10 µg/ml, respectively, in the presence and absence of 5 µM elacridar and 500 nM BafA1, and incubated for 2 h. YFP and GFP-tagged proteins were visualized using standard filter sets. All imaging was performed using a Zeiss Axio Observer Z1 microscope equipped with a digital camera (C10600/ORCA-R2 Hamamatsu Photonics). Images were taken using Zen software.

### Psychoactive Drug Screening Program

All information related to the PDSP can be found at https://pdsp.unc.edu/pdspweb/.

### Real Time-Quaking Induced Conversion (RT-QuIC) Assay

Recombinant bank vole PrP (residues 23 to 230; Methionine at position 109; accession no. AF367624) was expressed and purified as described previously (31). The concentration of recombinant PrP was determined using a molar extinction coefficient at 280 nm of 62,005/M/cm. RT-QuIC reactions were performed using black clear bottom 96-well plates and a reaction mixture of 10 mM phosphate buffer (pH 7.4), 300mM NaCl, 0.001% SDS, 1 mM EDTA, 10 µM ThT and 0.1 mg/ml PrP in a final volume of 98 µL. DMSO was limited to 0.02%. Reactions were seeded with 2 µL of the indicated dilution of RML infected or uninfected brain homogenate. Plates were sealed and incubated in a BMG FLUOstar plate reader at 42 °C with cycles of 1 min of shaking (700 rpm double orbital) and 1 min of rest. ThT fluorescence measurements were taken every 15 min (450 ± 10 nm excitation and 480 ± 10 nm emission; bottom read; 20 flashes per well; manual gain of 2000; 20 s integration time). Data was analyzed using GraphPad Prism.

### CRISPR/Cas9 mediated gene disruption

Knockout was achieved using kits from Synthego (Gene Knockout Kit v2), using three chemically modified sgRNAs for each locus (Rosa26 is targeted by a single guide): *Abcb1a* 5’-CAAAAUCGGAAUGUUCUUCC-3’, 5’-UAAUAGGAUUUACCCGUGGC-3’, 5’-CCAGCUGACAGUCCAAGAAC-3’; *Abcb1b* 5’-UCUAGUUUCGCUAUGCAGAU-3’,5’-AGAGGGGAAGUAAUGUUCCA-3’, 5’-AGUAAUGCUUGGCAGAAUAC-3’; *Abcg2*5’-GCGACAUUGGUACUAACACG-3’, 5’-UGCGAGCGUCCUAACGGCUC-3’, 5’-ACUCUUUACUUUCACUCGAU-3’; *Fech* 5’-UUUGAAUGGGAAGUGUCAUG-3’, 5’-UGAACUUCUCCAAGGGUUUC-3’, 5’-CUUGUGUAGGAAGCCAAAAA-3’. Recombinant Cas9 containing two nuclear localization signals (Synthego) is incubated with sgRNAs to form ribonucleoprotein complexes that are then electroporated into the cells using the Amaxa Cell Line Nucleofector kit V (Lonza Bioscience) with the manufacturer’s settings for N2a cells. To evaluate knockout efficiency, genomic DNA was extracted and targeted regions were amplified using Q5 high fidelity polymerase (New England Biolabs) using the following primers: *Abcb1a-*F 5’-ACTAGTCATGCCAAAGATACTAGG-3’, *Abcb1a*-R 5’-CCAACATAGGGCTGGCCTAG-3’, *Abcb1b*-F 5’-GCACCATGCTTTGAACACA-3’, *Abcb1b*-R 5’-TCATTACAAACCTGAATGAGAGCG-3’, *Abcg2*-F 5’-CATTACCGGCTGGGCTCAC-3’, *Abcg2*-R 5’-CAAGAGATCAAGTAACTTATGTCCTAGT-3’, *Fech*-F 5’-GGAGAGTGCTGGCTAACTGG-3’, *Fech*-R 5’-CCAACATAGGGCTGGCCTAG-3’. Sanger sequencing was performed with the following primers: *Abcb1a* 5’-GCCAAAGATACTAGGCAAAAATTAATGTAC-3’; *Abcb1b* 5’-CTTTGAACACAGCAGAGGGAGAG-3’; *Abcg2* 5’-CACAATCAAAGTGCTGGTATCTGTGTTGAT-3’; *Fech* 5’-CTGTAGGGCTCTGCATACCATTTC-3’; Editing efficiency was determined with the Synthego Inference of CRISPR Edits (ICE) tool (https://ice.synthego.com) (103) and by western blot. Oligonucleotide sequences for *Sigmar1*, *Tmem97*, and *Rosa26* targeting have been described (14).

### Primary Astrocyte Isolation, infection, and treatment

Astrocytes were isolated from neonatal mouse cerebella as described (104). Briefly, cerebella were isolated from p0 pups and the meninges were removed. Dissociation was performed using 2.5% trypsin and 1% DNAse I in 1x HBSS without calcium or magnesium. After incubation at 37°C, cells were spun down at 100 x g for 10 min. HBSS was removed and cells were resuspended in DMEM/F12 (Gibco) with 1% N2 supplement, 10% FBS, and 1% pen/strep and three cerebella were plated per flask into T75 flasks. Media was replaced 24 hours after isolation and allowed to grow for 48 hours before removing OPCs by tapping the flask. For prion infection, astrocytes were seeded into 24-well plates at 100,000 cells/well and allowed to grow overnight. The next day, they were exposed to 0.5% brain homogenate from terminally ill 22L mice. After 24 hours, the medium was replaced and cells were passaged after 2 and 10 days with medium changes every 2-3 days. Astrocytes from *Prnp^−/−^* mice (105) were infected in parallel to monitor the levels of residual inoculum. After the second passage, one set of cells (proliferative) were exposed to increasing concentrations of elacridar for 5 days (0.1% DMSO). Another set of cells was allowed to grow for 18 days following the second passage to induce contact inhibition of growth. Once deemed non-proliferative by eye, these cells underwent the same treatment regimen.

### RNA-seq

Total RNA was extracted using the RNeasy mini plus extraction kit (Qiagen) following manufacturer’s instructions. RNA purity and concentration were measured using a nanodrop (Thermo). Library preparation, sequencing, and differential gene expression analysis was performed by Innomics (Beijing, China). Gene set enrichment analysis (GSEA) was performed using the fgsea package in R using the Molecular Signatures Database (MSigDB) from the Broad Institute.

### LipidTox assay

Uninfected N2a cells were seeded in black 96-well glass bottom plates and incubated with 10 μM of compound or DMSO alone with 1x HCS LipidTOX™ Red neutral lipid stain (Invitrogen) for 24 hours. Cells were washed with 3x with PBS and fixed with 4% paraformaldehyde with 10 μg/ml Hoechest 33342 (Tocris) for 30 min at room temperature. After 3x washes with PBS, fluorescence was measured with a plate reader: ex/em 577/609 nm for LipidTox; 350/450 nm for Hoechest 33342. LipidTox signal was normalized to Hoechest 33342 signal for plotting. Images were taken with a Zeiss Axio Observer Z1 microscope using a 10x objective and Zen software.

### α-synuclein and tau seeding assays

TgM83^+/−^, Tg2541^+/+^, and C57BL/6J brain homogenates were subjected to PTA precipitation, as described (58). Assays were performed in a black 96-well clear-bottomed plate. HEK293T cells expressing α-syn140*A53T– YFP (58) or TauRD(LM)–YFP (60) were plated at 14,000 (α-syn) or 24,000 (tau) cells/well with 5 µM elacridar and allowed to attach for 3 hours. PTA precipitates were diluted 1:4 in PBS and incubated with Lipofectamine 2000 for 2 hours at room temperature. These mixtures were then diluted 4x with warm media and added to the cells and incubated for 4 days before imaging.

## Ethics statement

All experiments involving animals were conducted according to the United States Department of Agriculture Animal Welfare Act and the National Institutes of Health Policy on Humane Care and Use of Laboratory Animals. Ethical approval (AN-14997) was obtained from Boston University Chobanian & Avedisian School of Medicine institutional animal care and use committee

## Author Contributions

Conducted experiments: RCCM, NTTL, NMR, EF, GL; Designed experiments: RCCM, DAH; Contributed materials/methods: RCCM, NTTL, NMR, EF, GL, ICO, JRPG, DGF, AS, JSV, ABB, RC, GCT, DAH; Project supervision: RCCM, ABB, RC, GCT, DAH; writing/editing: RCCM, DAH.

## Acknowledgements

This work was supported by the National Institutes of Health grant number 5R01NS065244, awarded to DAH. RCCM was supported by grants from the Department of Defense (W81XWH-21-1-0141) and the Creutzfeldt−Jakob Disease Foundation. We are grateful to Amanda L. Woerman for the gift of α-syn140*A53T– YFP and TauRD(LM)–YFP HEK293T cells, and TgM83^+/−^ and Tg2541^+/+^ brain homogenates. We also thank Byron Caughey for supplying the plasmid to make the recombinant bank vole PrP used in RT-QuIC assays, Joel C. Watts and Ina Vorberg for cell lines used in this study, Eric Vallabh Minikel and Sonia M Vallabh for providing ZH3 *Prnp^−/−^* mice, and *Matthew Au for assistance with measurements of elacridar fluorescence.* K_i_ determination was performed by the National Institute of Mental Health’s Psychoactive Drug Screening Program, Contract # HHSN-271-2018-00023-C (NIMH PDSP). The NIMH PDSP is directed by Bryan L. Roth at the University of North Carolina at Chapel Hill and Project Officer Jamie Driscoll at NIMH, Bethesda MD, USA.

## Notes

### Competing Interest Statement

The authors have declared no competing interest.

### Summary of Updates

The Author list has been corrected and supplemental files moved from the main body

