## Supplementary Figures for "Lysosomal Enhancement Prevents Infection with PrP^Sc^, α-Synuclein & Tau Prions"

**Supplementary Figure 1: Uncropped western blots quantified in Figure 1A**

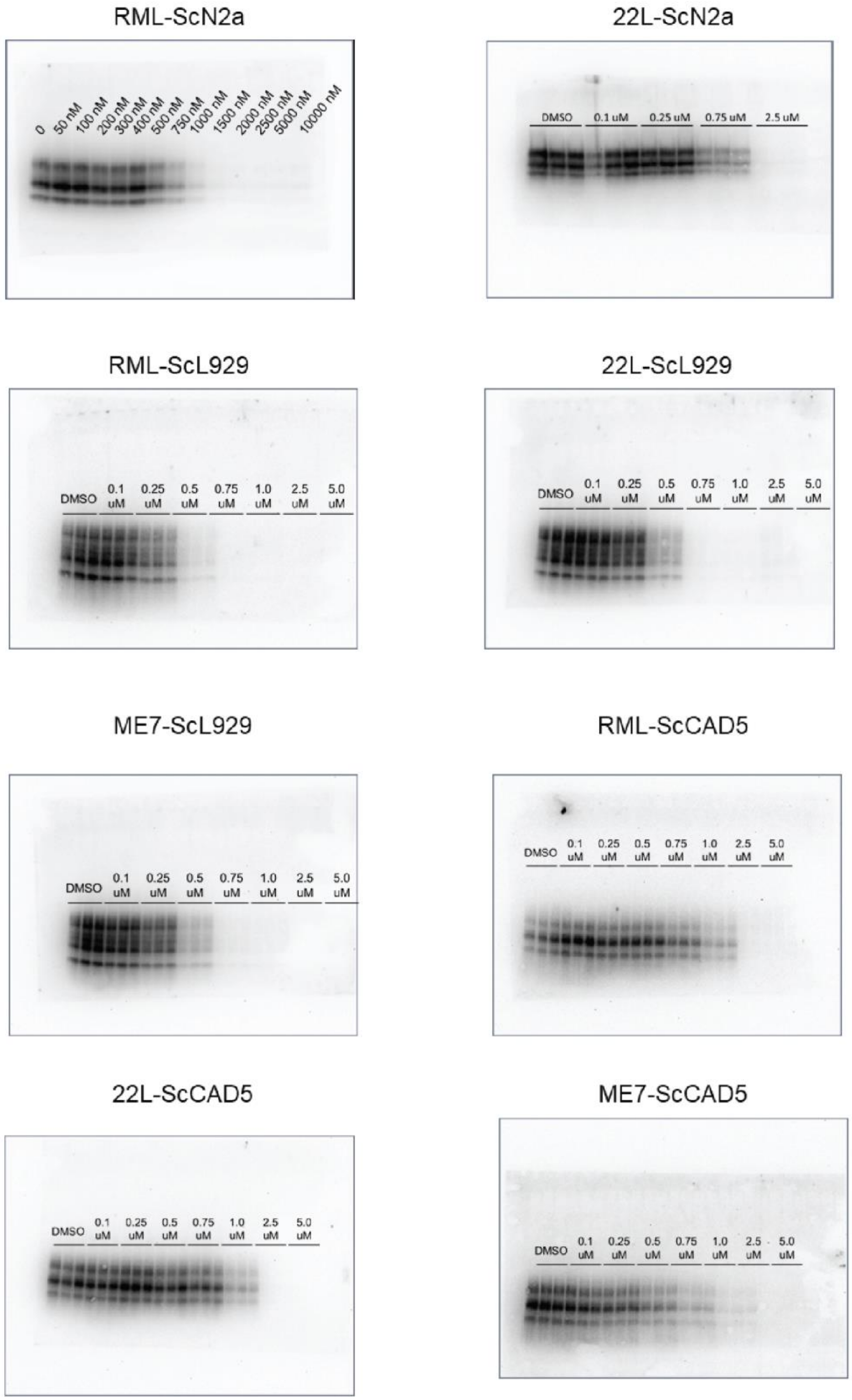

**Supplementary Figure 2: Elacridar is effective against mouse and sheep prions in RK13 cells**

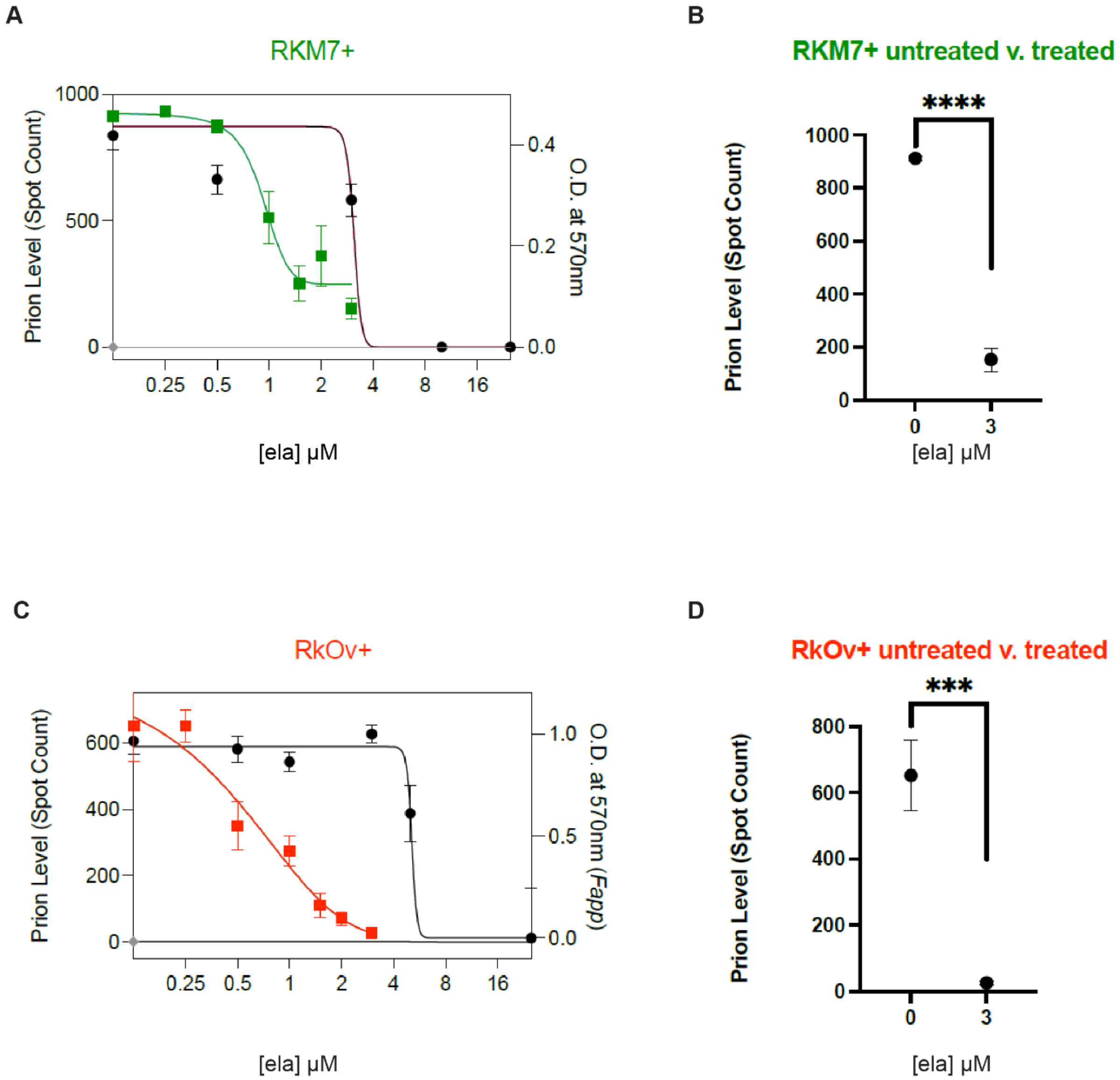


**A)** RKM7+ cells expressing murine $PrP^C$ were incubated with the indicated concentration of elacridar for 7 days before performing the scrapie cell and MTT assays in parallel. **B)** Direct comparison of spot counts from treated and untreated RKM7+ cells. **C)** RkOv+ cells expressing ovine $PrP^C$ were incubated with the indicated concentration of elacridar for 7 days before performing the scrapie cell and MTT assays in parallel. **D)** Direct comparison of spot counts from treated and untreated RkOv+ cells. For all quantification, $p < 0.0001$ = ****; $p < 0.001$ = ***; $p < 0.01$ = **; $p < 0.05$ = *; ns = not significant using an unpaired Student's t-test.

**Supplementary Figure 3: Elacridar is intrinsically fluorescent**

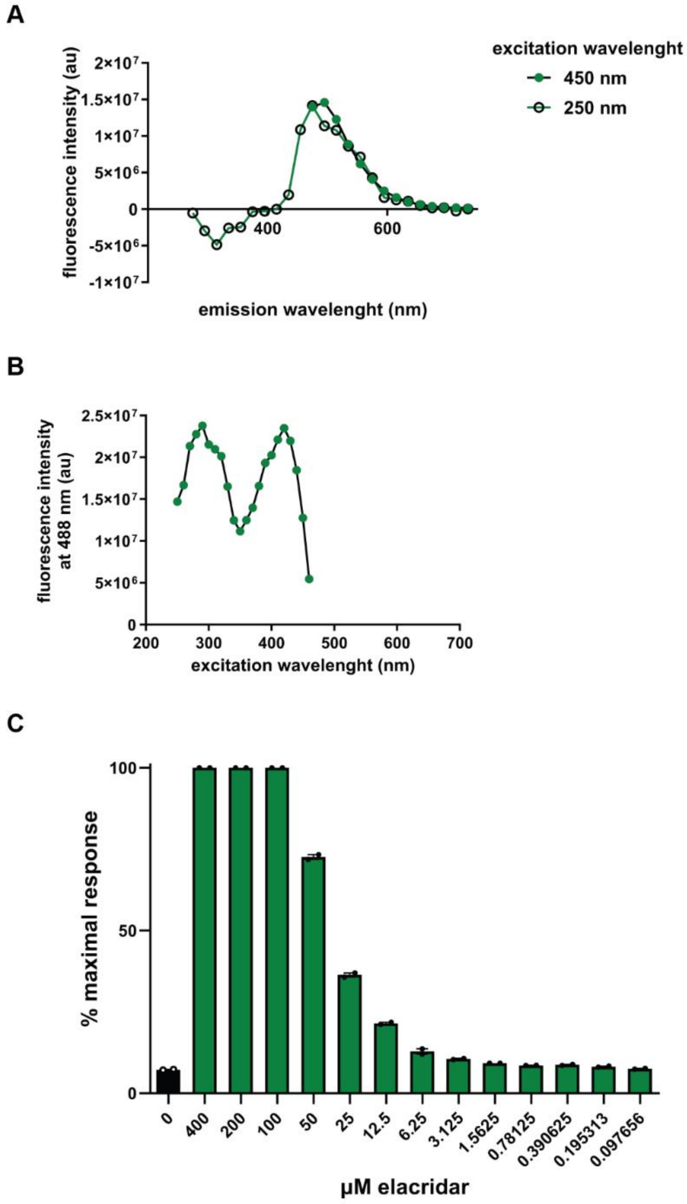


**A)** A 200 µM solution of elacridar in 1% DMSO was excited at 450 or 250 nm and emission spectra was recorded in 20 nm increments using a Tecan Infinite M1000 microplate reader. **B)** A 200 µM solution of elacridar in 1% DMSO was excited in 10 nm increments from 250-460 nm. Emission at 488 nm was recorded using a Tecan Infinite M1000 microplate reader. **C)** 100 µl of elacridar at the indicated concentration in RT-QuIC buffer was placed in black 96-well clear-bottomed plates and read in the BMG Polarstar plate reader at an excitation wavelength of 450 nm and emission wavelength of 480 nm. Percent maximal response is plotted, showing that the baseline fluorescence of elacridar begins increasing above ~5 µM. In RT-QuIC assays, this interferes with the ThT fluorescence read-out.

**Supplementary Figure 4: CRISPR/Cas9 knockout analysis and western blots for Figure 2F**

**A**

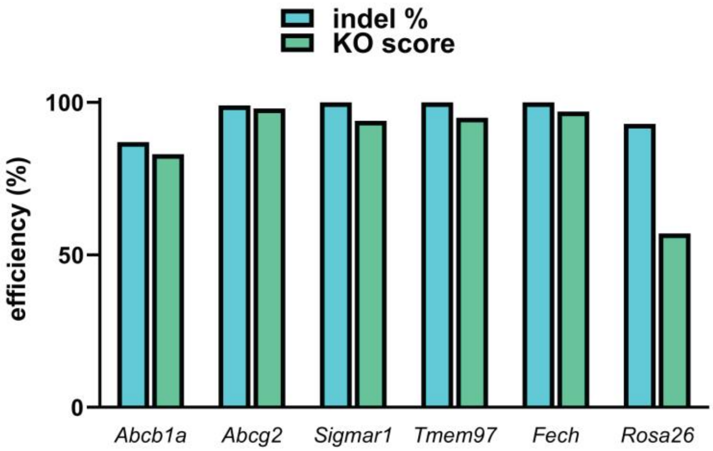


**B**

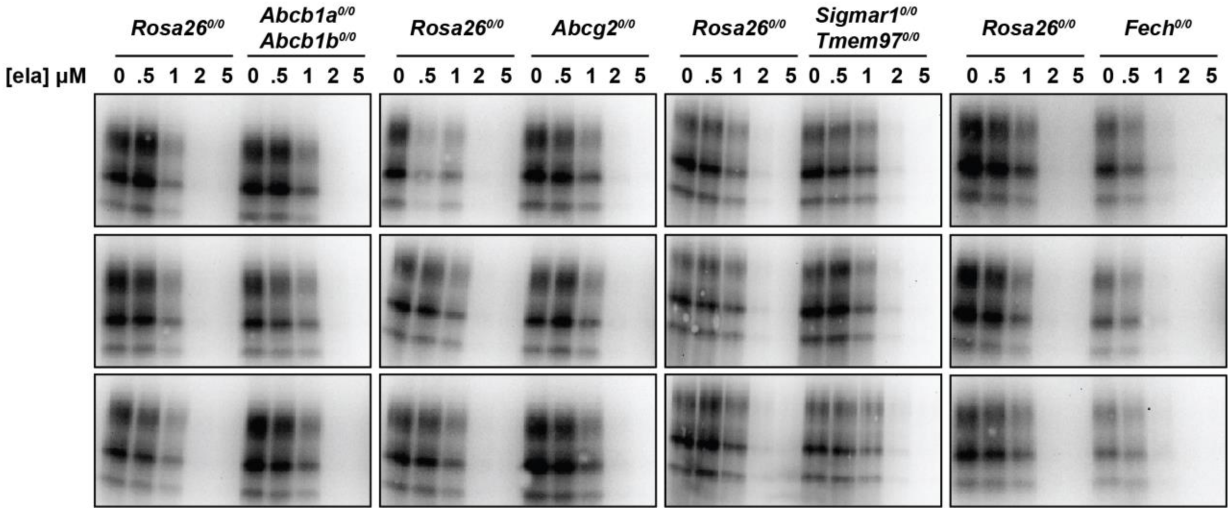


**A)** Inference of CRISPR Edits (ICE) analysis of *Abcb1a, Abcb1b, Abcg2, Sigmar1, Tmem97,* and *Fech* gene disruption. Following PCR amplification of targeted loci, ICE analysis allows for the deconvolution of Sanger sequencing data to provide indel % (editing efficiency) and a knockout score (KO: proportion of edits that result in a frame shift). The Rosa26 safe harbor locus is targeted with a single guide RNA as a control. **B)** Western blots of PK-resistant PrP corresponding to the graph shown in Figure 2F.

**Supplementary Figure 5: Western blots quantified in Figure 4**

**A**

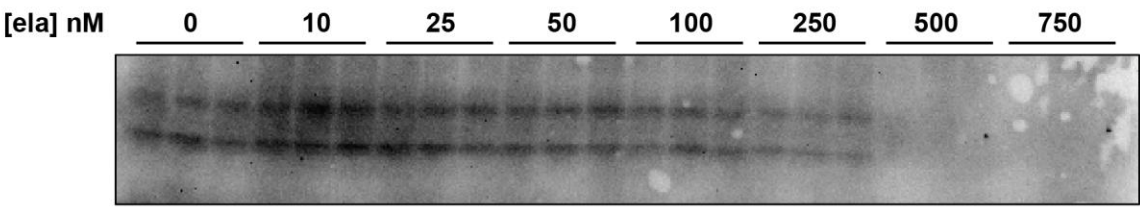


**B**

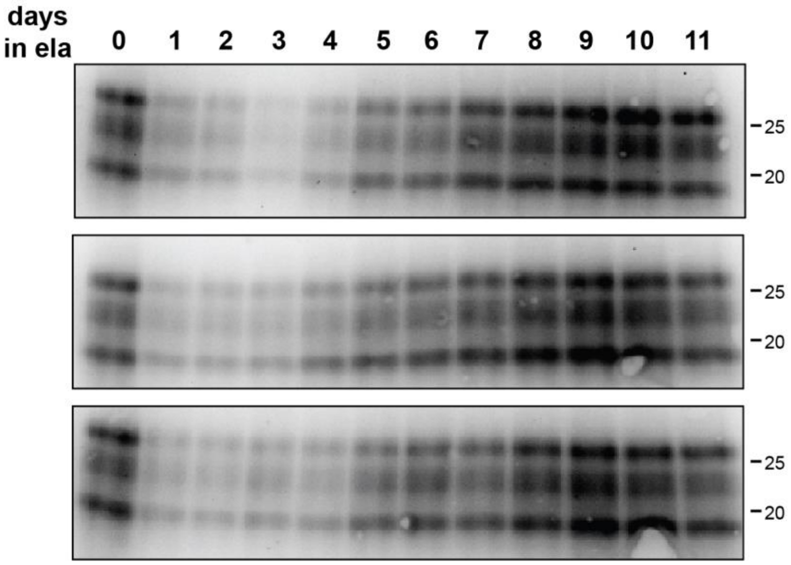


**A)** Western blots of PK-resistant PrP corresponding to the graph shown in Figure 4A. **B)** Western blots of PK-resistant PrP corresponding to the graph shown in Figure 4B.

**Supplementary Figure 6: Low magnification images of PIPLC treated N2a cells**

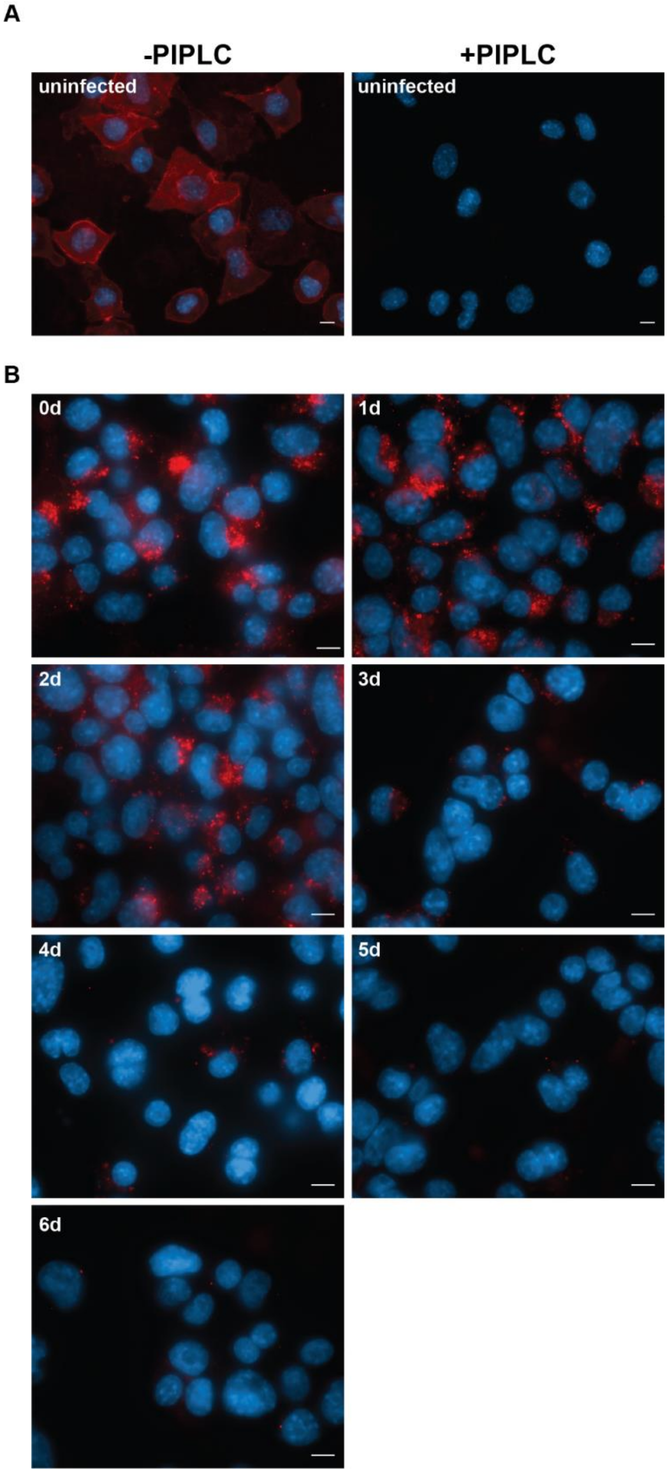


**A)** PrP immunofluorescent staining with D18 of uninfected N2a cells treated without (left) or with PIPLC (right). **B)** Low magnification images of RML-infected N2a cells treated with 2.5 µM elacridar for the indicated number of days before treatment with PIPLC/GdnHCl and imaging (corresponding to Figure 4C and D). Scale bars = 10 µm.

**Supplementary Figure 7: Lysosomal accumulation of elacridar is independent of known molecular targets**

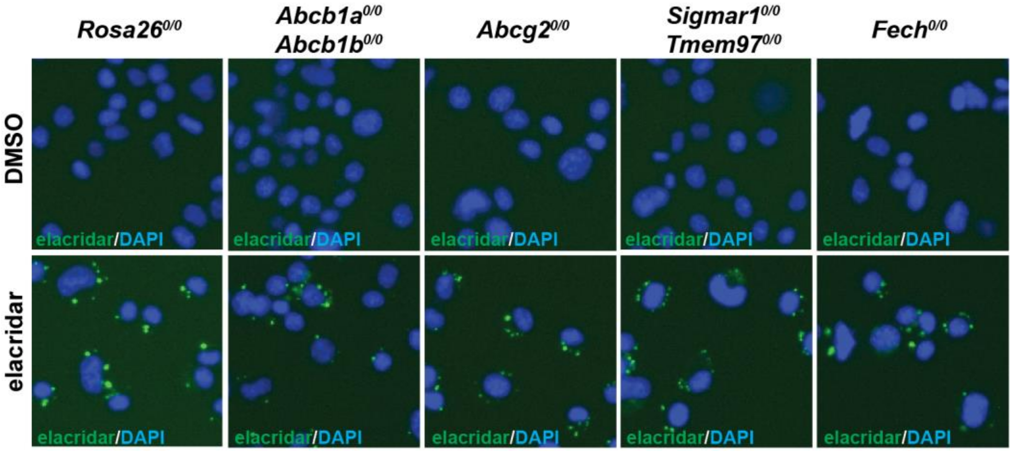


CRISPR-edited cells knocked out for expression of the indicated genes were incubated with 2.5 µM elacridar overnight before imaging.

**Supplementary Figure 8: Treatment with elacridar induces autophagy**

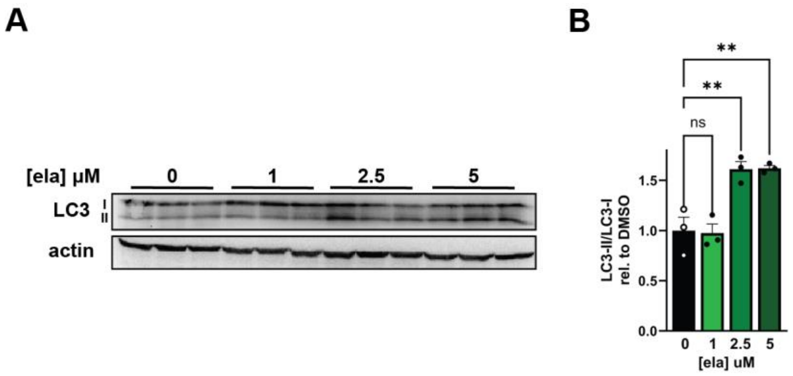


**A)** N2a cells were treated with increasing concentrations of elacridar before lysis and analysis of LC3 levels by western blot. **B)** Quantification of **A**. $p < 0.0001$ = ****; $p < 0.001$ = ***; $p < 0.01$ = **; $p < 0.05$ = *; ns = not significant using one-way ANOVA and Dunnett's multiple comparisons test.

**Supplementary Figure 9: Ammonium chloride prevents the lysosomal accumulation of elacridar**

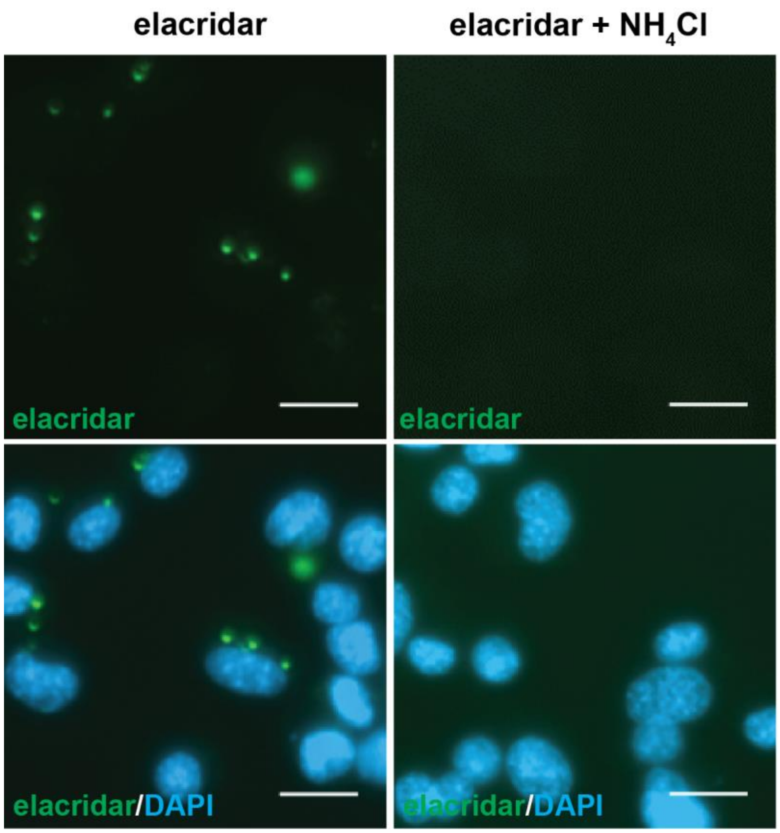


Cells were treated with 5 µM elacridar in the presence and absence of 10 mM $NH_4Cl$. Accumulation of elacridar in lysosomes is completely prevented by $NH_4Cl$.

**Supplementary Figure 10: LTR99 and DQ-BSA staining of N2a cells treated with quinacrine**

**A**

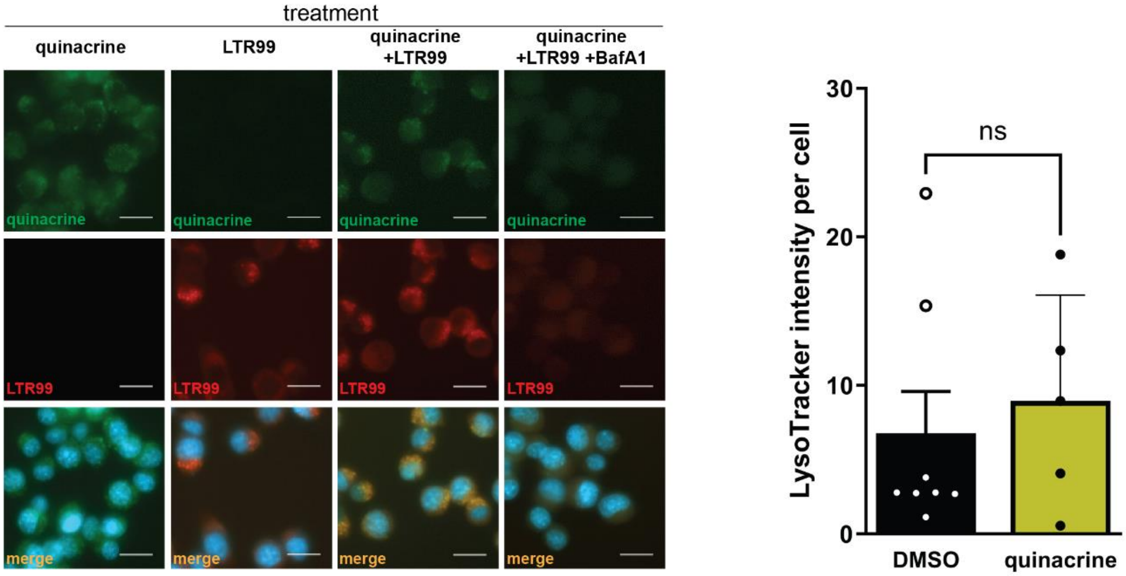


**B**

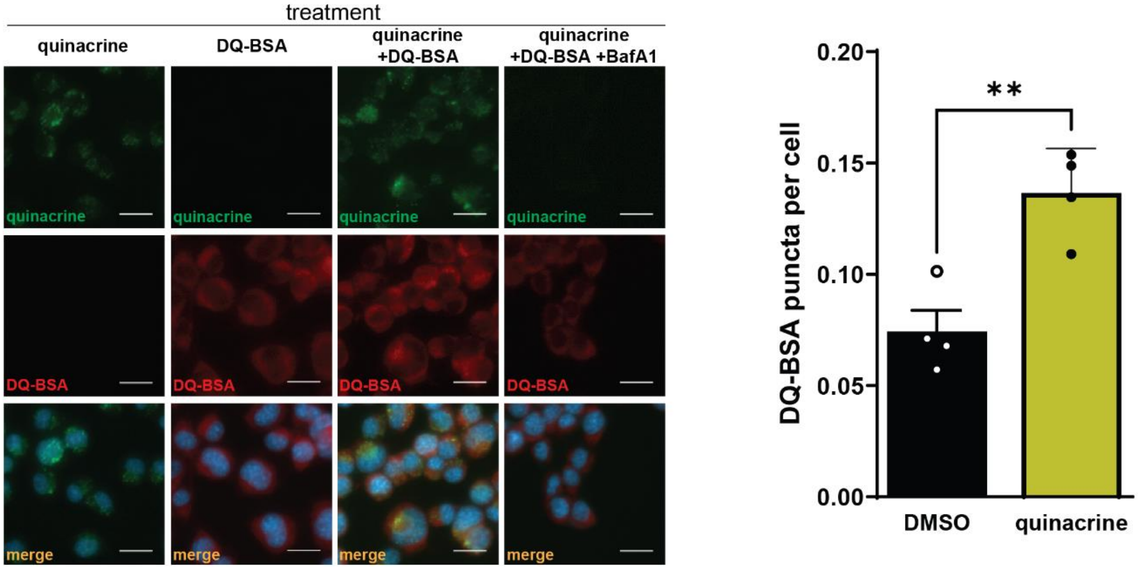


**A)** Quinacrine itself does not form punctate intracellular accumulations in N2a cells but co-localizes with LTR99 without increasing its fluorescence intensity. **B)** The application of quinacrine to N2a cells has a minor effect on DQ-BSA fluorescence. For all quantification, $p < 0.0001$ = ****; $p < 0.001$ = ***; $p < 0.01$ = **; $p < 0.05$ = *; ns = not significant using Student's t-test with Welch's correction.

**Supplementary Figure 11: Graphical abstract**

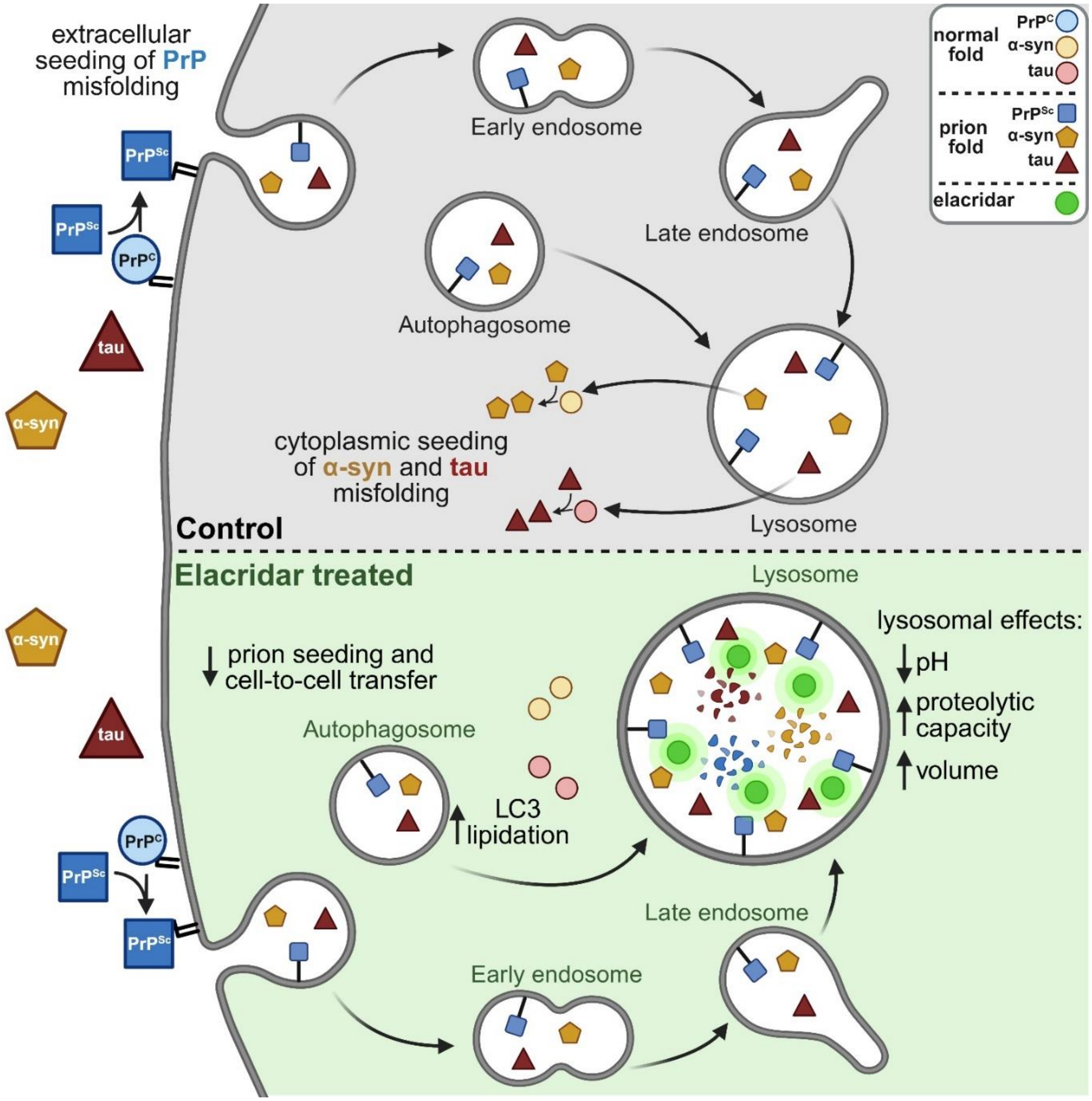
